# Origin and diversification of a globally distributed group of parasitic feather lice

**DOI:** 10.1101/2024.01.11.575230

**Authors:** Andrew D. Sweet, Jorge Doña, Kevin P. Johnson

## Abstract

Despite their extensive diversity and ecological importance, the history of diversification for most groups of parasitic organisms remains relatively understudied. Elucidating broad macroevolutionary patterns of parasites is challenging, often limited by the availability of samples, genetic resources, and knowledge about ecological relationships with their hosts. In this study, we explore the macroevolutionary history of parasites by focusing on parasitic body lice from doves. Building on extensive knowledge of ecological relationships and previous phylogenomic studies of their avian hosts, we tested specific questions about the evolutionary origins of the body lice of doves, leveraging whole genome data sets for phylogenomics. Specifically, we sequenced whole genomes from 68 samples of dove body lice, including representatives of all body louse genera from 51 host taxa. From these data, we assembled >2,300 nuclear genes to estimate dated phylogenetic relationships among body lice and several outgroup taxa. The resulting phylogeny of body lice was well supported, although some branches had conflicting signal across the genome. We then reconstructed ancestral biogeographic ranges of body lice and compared the body louse phylogeny to phylogeny of doves, and also to a previously published phylogeny of the wing lice of doves. Divergence estimates placed the origin of body lice in the late Oligocene. Body lice likely originated in Australasia and dispersed with their hosts during the early Miocene, with subsequent codivergence and host switching throughout the world. Notably, this evolutionary history is very similar to that of dove wing lice, despite the stronger dispersal capabilities of wing lice compared to body lice. Our results highlight the central role of hosts in driving macroevolutionary patterns of their parasites over long periods of time.

## INTRODUCTION

Understanding the factors that drive diversification in a group of organisms is a core focus of evolutionary biology (Ricklefs 2010, Jezkova and Wiens 2017). The history of the diversification of parasites is especially relevant, given their remarkable diversity and role in ecological communities (Poulin and Morand 2000, Lafferty et al. 2006, Dobson et al. 2008, Morand 2015). Although some parasitic taxa have received considerable attention (e.g., monogeneans (Poulin 2002), haemosporidian blood parasites (Ricklefs et al. 2014), parasitoid wasps (Santos et al. 2019)), the evolutionary histories for most groups of parasites remain understudied. There are several reasons for this discrepancy in research output. Many parasites exhibit cryptic morphology, making it challenging to evaluate taxonomic diversity (Miura et al. 2006, Nadler and Pérez-Ponce de León 2011). Obtaining parasite samples often requires access to infected hosts, the availability of which can vary based on host species and parasite prevalence (Clayton and Walther 1997, Justine et al. 2012, Hay et al. 2020). To deeply understand the diversification history of a group of parasites, information about their host’s diversification is also necessary, which is lacking for many host-parasite systems (Cruaud et al. 2012, Wang et al. 2019).

Investigations into the diversification and evolutionary history of parasites should address several primary objectives. First, studies of parasite diversification should establish when and where a group of parasites originated on their hosts (Wilson et al. 2012, de Vienne et al. 2013). Did a group of parasites have a single origin on a group of hosts or did they originate multiple times independently? Did multiple origins of parasites occur on hosts from different geographic regions? Addressing these questions ultimately relies on understanding the ecology and life history of hosts and their parasites (Clayton and Johnson 2003, Penczykowski et al. 2015, Clayton et al. 2016, Hembry and Weber 2020). For example, generalist parasites, and/or those with high dispersal capabilities, might often switch between various host groups (Vogwill et al. 2008, Sweet and Johnson 2018, Park et al. 2020). Assessing when parasites first formed associations with their hosts relies on dated phylogenies for both parasite and host. Estimating node ages can be challenging, as it requires fossil information, secondary calibrations, or reliable molecular clocks (Yang and Rannala 2006, Ho and Duchêne 2014). Many organisms lack this information, particularly groups of smaller-bodied parasites that often have poor fossil records (Koch 1978, Smith and Peterson 2002). Investigating where parasites originated on a group of hosts requires comparing historical biogeography of hosts and parasites, which again typically uses dated phylogenies. Historical biogeography can also be sensitive to sampling bias; thus, a comprehensive representation of both host and parasite taxa is crucial. Sampling can prove challenging for both host and parasites, especially for rare hosts or hosts from isolated geographic locations.

Another important question to address in studies of parasite evolution is how much a host influences the diversification history of a parasite (McCoy et al. 2003, Barrett et al. 2008, Michelet and Dauga 2012). Obligate parasites are expected to have an evolutionary history mirroring that of their hosts (i.e., Fahrenholz’s Rule), implying that the phylogeny and historical biogeography of a host will predict their parasites’ phylogenetic and biogeographic history (Fahrenholz 1913, Eichler 1948, Timm 1983). In reality, very few host-parasite systems strictly follow the expectations of Fahrenholz’s Rule, even for parasites with obligate, host-specific relationships with their hosts (Paterson et al. 2000, Benovics et al. 2019). However, hosts can still play a major role in the diversification of their parasites (Hafner and Nadler 1988, Hughes et al. 2007). The effects of host-driven evolution can manifest in the phylogenetic and biogeographic patterns of parasites (Weckstein 2004, Blasco-Costa et al. 2021). Thus, a broad macroevolutionary approach at various timescales is necessary to fully understand the role of hosts in shaping the evolutionary history for a group of parasites (Jackson et al. 2008, Sweet et al. 2016).

Host organisms also often harbor multiple groups of similar parasites, which can further complicate, but also inform, studies of parasite diversification (Bordes and Morand 2009). For example, different groups of parasites compete for resources and space on a host, influencing their macroevolutionary trajectories (Penczykowski et al. 2015, Harmon et al. 2019, Hembry and Weber 2020, Dismukes et al. 2022). Thus, focusing on the evolutionary history of only a single group of parasites could be missing an important driver of diversification, i.e. other similar parasites. Comparing multiple groups of similar parasites from a single group of hosts can allow for testing predictions about comparative evolution between hosts and parasites, especially if the parasites have variation in life history or ecology (Weiblen and Bush 2002, Clayton and Johnson 2003). For example, if one group of parasites has relatively strong dispersal capabilities or can outcompete other similar parasites, those parasites are likely better able to establish populations on different host species (i.e., host switching) (Hoberg and Brooks 2008, Ellis et al. 2015). Frequent host switching can result in considerable incongruence between host and parasite evolutionary trees (Paterson and Banks 2001). However, the ecology of a group of parasites needs to be well-understood to make a link between the process of dispersal and host switching.

Parasitic body lice from pigeons and doves (“doves” hereafter) are an excellent system for studying the diversification of parasites. These lice are broadly distributed and the ecology of both the lice (Insecta: Phthiraptera) and their dove (Aves: Columbiformes) hosts is relatively well-studied (Goodwin 1983, Gibbs et al. 2001, Clayton et al. 2016). The body lice of doves are currently placed in five genera (*Auricotes, Campanolutes, Coloceras, Kodocephalum,* and *Physconelloides*) (Price et al. 2003), although have been variously placed in additional genera by some authors. These lice are permanent and obligate ectoparasites that consume feathers and typically complete their approximately one-month lifecycle on a single host individual (Nelson and Murray 1971, Marshall 1981). Adult females lay their eggs by cementing them to feather barbs, and juveniles complete three nymphal instars before reaching adulthood. Members of these genera are termed “body” lice because they escape host preening defenses by burrowing in the downy body feathers and have a distinctive rounded body form and rounded head margin distinguishing them from other ecomorphs (Clay 1949). Body lice are suspected to occur on nearly all of the ∼350 species and 49 genera of extant doves, given that most species that have been extensively examined for ectoparasites have been shown to harbor body lice (Price et al. 2003). As with their hosts, body lice are globally distributed, inhabiting every continent except Antarctica (Gibbs et al. 2001).

Past phylogenetic work has helped establish some knowledge about the evolutionary history of both doves and their body lice. For body lice, previous studies based on Sanger sequencing data sets of just two or three mitochondrial and nuclear genes (Johnson et al. 2001, 2011; Sweet et al. 2017), while resolving some relationships, produced trees with many poorly supported branches. The prior study with the most extensive taxon sample (Johnson et al. 2011), comprising 71 samples of dove body lice, found evidence that dove body lice originated from a host-switch from landfowl (pheasants, quail, partridges, grouse, megapodes, etc.; Aves: Galliformes) to doves (Columbiformes). The tree in Johnson et al. (2011) also indicated that there may have been an additional host-switch back to landfowl, but generally inter-ordinal switches in this group were inferred to be very rare. The original host switch from landfowl to doves could have facilitated an increase in diversification of dove body lice as they radiated across a group of hosts (Hay et al. 2020). These previous studies also failed to recover any of the five body louse genera as monophyletic, but the backbones of these trees were not well supported. Regarding the dove hosts, although there have been many prior Sanger-based sequencing studies (Johnson and Clayton 2000, Pereira et al. 2007, Johnson et al. 2010, Johnson and Weckstein 2011, Cibois et al. 2014, Sweet and Johnson 2015, Nowak et al. 2019), these generally did not result in well supported trees, especially along the backbone. In contrast, Boyd et al. (2022) used whole genome sequence data and fossil information to estimate a well-supported and dated phylogenetic tree of 61 species of doves. They also conducted historical biogeographic analyses and found support for a New World/Australasian origin of doves before the breakup of Gondawana (∼50-60 MYA). This study provided evidence for widespread long-distance dispersal of doves from Australasia and the New World during the early Miocene.

In addition to hosting body lice, doves harbor another group of feather lice, “wing lice” in the genus *Columbicola*. Wing lice are not closely related to dove body lice, with *Columbicola* being on a long branch that is sister to most other true feather lice in the Parvorder Ischnocera (Johnson et al. 2018, de Moya et al. 2021). Dove wing lice also differ from dove body lice in the ecological relationships with their hosts. Wing lice escape from host preening defenses by inserting between the feather barbs of their host’s wing feathers (Clayton et al. 2003) and have a long and slender form adapted for this behavior. Like body lice, wing lice eat the downy portions of the hosts body feathers (Clayton 1991). However, body lice are more likely to outcompete wing lice without mediation from host anti-parasite behavior (Bush and Malenke 2008). Wing lice are also better able to disperse among different hosts compared to body lice. Wing lice are known to hitchhike on winged hippoboscid flies, whereas body lice do not engage in this phoretic behavior (Harbison et al. 2009). These differences in dispersal ability have implications for patterns of codiversification between doves and their lice, with body lice expected to have stronger congruence with their hosts at phylogenetic and population genetic scales (Clayton and Johnson 2003, DiBlasi et al. 2018, Sweet and Johnson 2018). However, cophylogenetic patterns can vary at different evolutionary time scales (Sweet et al. 2016, 2018). A well-supported phylogenomic tree from 61 taxa of *Columbicola* (Boyd et al. 2017) indicated a New World + Australasian origin for *Columbicola* with subsequent dispersal events to other continents, which is consistent with the evolutionary history of their dove hosts. Comparisons of the phylogeny of *Columbicola* to the phylogeny of their dove hosts (Boyd et al. 2022) showed evidence of both cospeciation and host switching, with a relative increase in cospeciation over time. Thus, it would be interesting to compare the codiversification of body lice to these patterns in wing lice.

Here, we used genomic data from a worldwide sample of dove body lice to better understand the diversification of this group of parasites. More specifically, our goals were to estimate a well-supported and dated phylogeny of dove body lice, reconstruct the historical biogeography of the group, compare the phylogenies of body lice and their dove hosts, and compare the patterns of diversification between dove body lice and wing lice. Using these approaches, we addressed three questions related to the diversification of body lice: 1) When and where did body lice originate on their hosts? 2) How have body lice diversified with their hosts over time, and 3) How does the diversification history of body lice compare to that of dove wing lice? Our results not only enable us to address these specific questions related to the evolutionary history of lice, but also provide broad insight into how parasites interact with their hosts on a macroevolutionary scale.

## METHODS

### DNA extraction and sequencing

Taxon sampling included 68 samples of dove (Columbiformes) body lice that had been stored in 95% ethanol at -80°C. The samples included lice from every continent on which doves occur, representing five genera of dove lice from 51 different species of doves. We also included samples of three genera of lice from 12 species of landfowl (Galliformes), and 12 outgroup taxa representing major clades of Ischnocera (Supplementary Table S1). Obtaining a sufficient quantity of DNA for genomic sequencing required an extraction process that destroyed the louse specimens, so it was not possible to retain specimens as slide vouchers. However, before extraction, each louse specimen was photographed as a voucher (images available in Dryad).

Each louse specimen was identified to genus and species (when possible) using existing taxonomic keys and/or Price et al. (2003). These identifications were also supplemented by comparison of the assembled mitochondrial cytochrome c oxidase I (*cox1*) sequences (see below) to previously published Sanger sequences for this gene in which a slide mounted specimen was prepared. We extracted genomic DNA (gDNA) from individual lice (i.e., not pooled individuals) using a modified protocol with Qiagen QIAamp DNA Micro Kits (Qiagen, Hilden, Germany). Specifically, we incubated each specimen in a proteinase K and buffer solution for ∼72 hours to maximize DNA concentration in the final elution. We then quantified each extraction using a Qubit fluorometer. From these total gDNA extractions, we prepared libraries using the Hyper Library construction kit (Kapa Biosystems) tagging them with unique dual-end adaptors. The libraries were sequenced using the Illumina NovaSeq 6000 platform with S4 reagents, and libraries were pooled 48 to one lane and sequenced for 150 bp paired-end reads. This multiplexing was estimated to achieve at least 30-60X coverage of the nuclear genome for each sample, assuming a ∼200 Mbp genome size.

### Sequence quality control and assembly

We ran quality checks on the resulting paired-end read files using FastQC (Babraham Bioinformatics) to check for poor quality reads, adapter content, and high levels of sequence duplication. We then used fastp v0.20.1 (Chen et al. 2018) to perform adaptor and quality trimming (phred quality >= 30).

For the assembly of nuclear loci, we initially prepared trimmed libraries and converted them into aTRAM 2.0 (Allen et al. 2018) blast databases, using the atram_preprocessor.py command from aTRAM v2.3.4. Our reference set consisted of 2395 single-copy ortholog protein-coding genes derived from the human louse, *Pediculus humanus*, a set previously employed in phylogenomic studies on lice (Johnson et al. 2018, 2021, 2022). We ran aTRAM assemblies (using atram.py command) using tblastn, on the amino acid sequences of the reference genes, and then used the ABySS assembler (iterations = 3, max-target-seqs = 3000) (Simpson et al. 2009). We then stitched together the exon sequences from these protein-coding genes using the Exonerate (Slater and Birney 2005) pipeline within aTRAM (the atram_stitcher.py).

To assemble mitochondrial cytochrome c oxidase I (*cox1*) gene, we took into account the high coverage of mitochondrial sequence reads within these Illumina raw read datasets, often surpassing 1000X. To manage this, we used Seqtk v 1.3 (https://github.com/lh3/seqtk) to subsample a total of four million reads (two million each of read1 and read2) per library (Shen et al. 2016). This step aimed to avoid assembly errors or potential contaminants. We used a previously published complete *cox1* sequence from *Campanulotes compar* (Song et al. 2019) as a reference. We then used aTRAM 2.0 (Allen et al. 2018) for the assembly (1 iteration using ABySS).

### Phylogenetic inference

We first concatenated the DNA sequences from each sample for each gene using a custom R script. After this, we translated the nucleotide sequences to amino acids utilizing a custom Python script, followed by an alignment based on the amino acid sequences. For alignment, we used MAFFT v7.471 (--auto --preservecase --adjustdirection --amino) (Katoh et al. 2002, Katoh and Standley 2013). After alignment, we back-translated these aligned amino acid sequences to DNA sequences using the same Python script. We then filtered each gene alignment in trimal v.1.4 by removing sites with more than 60% missing data (“40% alignment”) and with a more stringent filter removing sites with more than 10% missing data (“90% alignment”) (Capella-Gutiérrez et al. 2009). For each dataset (full, 40%, and 90%), we concatenated the gene alignments using AMAS (Borowiec 2016). We also tested for evidence of multiple substitution/saturation in the third codon positions. We separated third codon positions from first+second codon positions in Geneious Prime v.2023.2.1 and compared uncorrected vs. corrected (using the F81 substitution model) genetic distances in each alignment subset using the APE package v.5.7-1 (Paradis and Schliep 2019) in R v.4.3.1 (R Core Team 2023). Plots of genetic distances suggested the presence of multiple substitution in the third codon position, so we created additional data sets for analysis by removing third codon sites from the full, 40%, and 90% gene alignments and concatenated these filtered alignments with AMAS.

We estimated phylogenetic relationships among our taxa using both concatenated and coalescent-based approaches for all six datasets (all sites and 1+2 sites for the full, 40%, and 90% alignments). For the concatenated alignments, we partitioned the alignments by gene and tested for optimal substitution models and partitions using ModelFinder based on Bayesian Information Criterion (BIC) with rcluster = 10, which considerably reduces computational time (Kalyaanamoorthy et al. 2017). We then ran a Maximum Likelihood (ML) analysis in IQ-TREE 2 v.2.0.3 (Minh et al. 2020b) using the optimal modeling scheme from ModelFinder. We assessed the resulting ML tree using 1,000 UltraFast bootstrap (BS) replicates (Hoang et al. 2018) and 1,000 RELL bootstrap replicates for the SH-like approximate likelihood ratio test (SH-aLRT) (Guindon et al. 2010). We used ASTRAL-III v.5.7.7 as a coalescent-based approach to estimate a species tree from gene trees (Zhang et al. 2018). We estimated gene trees in IQ-TREE 2 using ModelFinder to test for optimal substitution models for each gene. We then summarized the gene trees in ASTRAL and assessed branch support with local posterior probabilities (LPP) (Sayyari and Mirarab et al. 2016).

We ran several additional tests to assess our phylogenetic trees. First, we calculated gene concordance factors (gCF) and site concordance factors (sCF) for each of the ML trees generated from concatenated alignments (Minh et al. 2020a, Mo et al. 2023). The metrics gCF and sCF calculate the proportion of genes and sites (respectively) that support each split in a given phylogeny. We ran all gCF and sCF calculations in IQ-TREE2, using the gene trees, concatenated alignments, and concatenated trees for each of the six datasets. Second, we ran Approximately Unbiased (AU) tests to statistically compare ML phylogenies generated from the six different concatenated alignments (Shimodaira et al. 2002). We ran the AU tests in IQ-TREE 2 with 10,000 RELL bootstrap replicates and the 40%-complete concatenated dataset (sites with maximum 60% missing data) as the input alignment. Third, we compared all phylogenies, including from concatenated and ASTRAL analyses, using normalized Robinson-Foulds (RF) distances using the RF.dist command in the PHANGORN v.2.11.1 package in R (Robinson and Foulds 1981, Schliep 2011). Finally, to explore an unstable branch, we assessed quartet support for a branch in the ML phylogeny involving four subsets of taxa: 1) *Auricotes affinis*, 2) *Coloceras furcatum*, 3) a clade containing representatives of the genera *Campanulotes*, *Coloceras*, and *Physconelloides*, and 4) a clade containing representatives of the genera *Kodocephalum*, *Auricotes*, and *Campanulotes* (*Saussurites*). Our phylogenetic results supported these four subsets as major lineages, but the relationships among these lineages were not consistent or well-supported across all analyses. We assessed support for the relationship among the four subsets using four-cluster likelihood mapping in IQ-TREE 2 (Strimmer and von Haeseler 1997). We assigned the taxa to their respective clusters (i.e., subsets) and conducted likelihood mapping with 10,000 randomly drawn quartets.

### Divergence time estimation

We estimated divergence times for the lice using two different methods, both based on the ML topology derived from the 40% concatenated matrix. First, we estimated dates using MCMCTree in PAML v.4.9 (Yang 2007, dos Reis and Yang 2011). We estimated substitution rates using BASEML with a GTR model and a strict clock. We set the root age to 57 million years (MY) based on the upper 95% highest probability densities (HPD) divergence estimate of our root node from de Moya et al. (2021). We set the root age to <60 MY as a safe constraint and applied a GTR model with five gamma categories and alpha set to 0.365 (estimated from BASEML). For internal calibrations, we used secondary information from divergence estimates of seven terminal sister pairs of host species that likely codiverged with their lice (Supplementary Table S2). We used 95% HPD from the dating analysis for the host tree of Boyd et al. (2022) as soft bounds for each calibration. We first ran MCMCTree to estimate branch lengths, gradient, and Hessian. We then ran two independent MCMCTree runs to estimate divergence times, each with 500,000 Markov Chain Monte Carlo (MCMC) chain iterations (5,000 samples, sampling every 100 iterations) and 50,000 iterations discarded as a burnin. We assessed Effective Sample Sizes (ESS) of each run in Tracer v.1.7.2 (Rambaut et al. 2018) to ensure the chains reached stationarity (ESS >200). Second, we estimated ages using the least squares dating (LSD2) method from IQ-TREE 2 (To et al. 2016). We set the root age to 57.4 MY and set minimum and maximum node calibrations using the same seven 95% HPD divergence estimates from Boyd et al. (2022). We also resampled branch lengths 1,000 to generate confidence intervals and set minimum branch lengths to 0.1 to avoid having comb-like branches.

### Historical biogeography of dove body lice

We reconstructed the historical biogeography of dove body lice using the R package BioGeoBEARS v.1.1.3 (Matzke 2018). We used the time-calibrated phylogeny generated from the first MCMCTree run and trimmed the outgroup taxa. We also trimmed the phylogeny based on an Operational Taxonomic Unit (OTU) assessment using the *cox1* data. Following a similar approach to the one used for the nuclear loci, we translated the *cox1* DNA sequences to amino acids, aligned them (using MAFFT), and then back-translated them. We then estimated a phylogenetic tree using IQ-TREE 2 and calculated uncorrected genetic distances (p-distances) with the APE package in R. We then collapsed pairs of taxa with *cox1* distances <5% into a single terminal branch on the time-calibrated phylogeny by trimming one of the taxa with the drop.tip command in the APE package in R (Supplementary Figure S1). We then assigned biogeographic ranges to louse taxa according to the native ranges of their host species based on Gibbs et al. (2003) and Birds of the World (birdsoftheworld.org). We assigned each taxon to at least one of five biogeographic regions: Africa, Eurasia, Oceania/Australasia, North America, and South America. We assessed six different biogeographic models in BioGeoBEARS using weighted AIC corrected for small sample sizes (AICc): DIVA-like, DEC, BayArea-like, and each of these three models with the jump (J) parameter. Although the J parameter has been criticized (Ree and Sanmartín 2018), it is appropriate for the body louse system given the ability of their hosts to disperse through long-distance flight. For each model, we set the maximum number of areas to five.

### Cophylogenetic analysis

We compared the phylogenies of dove body lice and their dove hosts using the event-based approach implemented in Jane v.4.01 (Conow et al. 2010). Jane uses a Genetic Algorithm (GA) to reconcile two phylogenies with *a priori* costs assigned to explicit evolutionary events. Optimal reconciliations have the lowest overall costs. For the louse phylogeny, we used our ML tree estimated from the 40% concatenated alignment. We removed outgroup taxa, lice from galliform birds, and collapsed multiple representatives of a single OTU based on *cox1* distances (as detailed above). For the dove phylogeny, we used the ML topology from Boyd et al. (2022). However, not all host species were represented in the Boyd et al. (2022) phylogeny, so we used the R function bind.tip from PHYTOOLS v.1.2-0 (Revell, 2012) to incorporate these missing dove taxa into our tree based on other prior phylogenetic results. Specifically, we added *Geopelia striata* (Sweet et al. 2017), *Columba sjostedti* (Johnson 2004), *Claravis pretiosa*, *Columbina passerina, C. squammata, C. inca,* and *Uropelia campestris* (Sweet et al. 2015). We also removed outgroup taxa and tips that did not have an associated louse represented in the parasite phylogeny. We ran Jane on the edited phylogenies using default costs for each event (Cospeciation: 0, Duplication: 1, Duplication and Host Switch: 2, Loss: 1, Failure to diverge: 1), GA parameters set as number of generations = 100 and population size = 500, and 100 random tip mappings to assess whether the best cost is significantly lower than with random host-parasite associations. We also tested for the congruence between the dove louse and dove phylogenies using PACo, a distance-based approach that tests whether two phylogenies (as distance matrices) are independent of each other (Balbuena et al. 2013). Rejecting the null hypothesis indicates the two trees are significantly congruent (i.e., not independent). We used the modified dove tree from Boyd et al. (2022) and trimmed body louse tree, converted the trees to patristic distance matrices using the *cophenetic* command in R, and ran PACo for 999 iterations with the “r0” method (assuming parasites track their hosts) correcting for negative eigenvalues with the Cailliez correction (Cailliez 1983). We also calculated the link residuals, which indicates the contribution of specific host-parasite links to the overall phylogenetic congruence. We ran PACo in the PACO v.0.4.2 package in R (Hutchinson et al. 2017).

We also compared the dove body and wing louse phylogenies using PACo. We used the ML body louse tree estimated from the 40% concatenated alignment and wing louse tree from Boyd et al. (2022). We assigned associations between lice based on shared host species and removed louse taxa that did not share at least one host with the other group of lice. We converted the trimmed phylogenies to patristic distance matrices and ran PACo in the PACO R package with 999 permutations and the Cailliez correction for negative eigenvalues. We used the “swap” randomization algorithm, which maintains the number of interactions for every tip on both phylogenies, because interactions between the two groups of lice are likely not determined by one group or the other (i.e., wing lice are not dependent on body lice and vice versa). We also randomized the wing and body louse phylogenies 1,000 times to test whether the observed residual value is significantly less than a distribution of residuals from random pairs of trees.

## RESULTS

### Phylogeny

Whole genome sequencing (WGS) from our 80 ingroup samples of body lice produced an average of 106,166,506 reads per sample. Trimming poor quality and duplicate reads reduced the average number of reads to 96,766,486 per sample (Supplementary Table S1).

Assembly with aTRAM produced 2,367 orthologous genes. All three alignment data sets (unfiltered, 40% complete, 90% complete) had an average of 91 taxa out of 92 (80 ingroup and 12 outgroup taxa) across the 2,367 genes (1.1% missing). The average lengths of the gene alignments were 2,019 bp for the unfiltered alignments, 1,656 bp for the 40% complete aligments, and 1,611 bp for the 90% complete alignments. The concatenated alignments were a total of 4,779,222 bp for the unfiltered alignments (18.2% gaps), 3,919,395 bp for the 40% complete alignment (1.0% gaps), and 3,812,517 bp for the 90% complete alignment (0.2% gaps).

Concatenated and coalescent-based phylogenetic analyses both produced well-supported phylogenies of lice (Figure 1, Supplementary Figures S2-S6). All but three in-group branches received 100% bootstrap (BS)/SH-aLRT support and 1.0 local posterior probability (LPP) support with the full, 40% complete, and 90% complete alignments. Plots of corrected versus uncorrected distances suggested the third codon position has evidence of multiple substitution/saturation (Supplementary Figure S7). However, phylogenetic analyses of only the first and second codon positions did not improve branch support, and in some instances resulted in worse support (e.g., 82 vs. 93 BS for the clade of *Coloceras* spp. from *Macropygia*, *Reinwardtoena*, and *Chalcophaps indica*) (Supplementary Figures S8-S12). The topologies were very similar among all three data sets, including results from all codon position and only 1^st^+2^nd^ codon positions. Normalized RF distances were low across all pairwise comparisons of phylogenies, with many trees having RF=0 (average 0.04 across all pairs) (Supplementary Table S3). AU tests significantly rejected seven phylogenies as alternative topologies based on the 40% complete concatenated alignment, whereas the tests failed to reject three phylogenies (the three phylogenies from the concatenated alignments with all sites) as alternative topologies (Supplementary Table S4). The largest differences were between trees from the coalescent analysis of the 90% complete alignment with third codon positions removed and the unfiltered concatenated alignment with third codon positions removed (RF=0.09). Notably, all analyses recovered dove body lice as a monophyletic group and landfowl lice as paraphyletic, both with high support (100 BS/100 SH-aLRT/1.0 LPP for both branches). We also recovered *Coloceras furcatum* from *Lopholaimus antarcticus* as sister to the rest of dove body lice. Within dove body lice, all of the genera with multiple representatives were recovered as paraphyletic or polyphyletic. *Coloceras*, *Campanulotes*, and *Physconelloides* were recovered as polyphyletic, whereas *Auricotes* was paraphyletic with *Auricotes affinis* as sister to the rest of dove body lice excluding *Coloceras furcatum*. *Physconelloides* was separated into two clades, one from New World taxa and one from Australian taxa, separated by *Campanulotes frenatus*.

**Figure 1.**
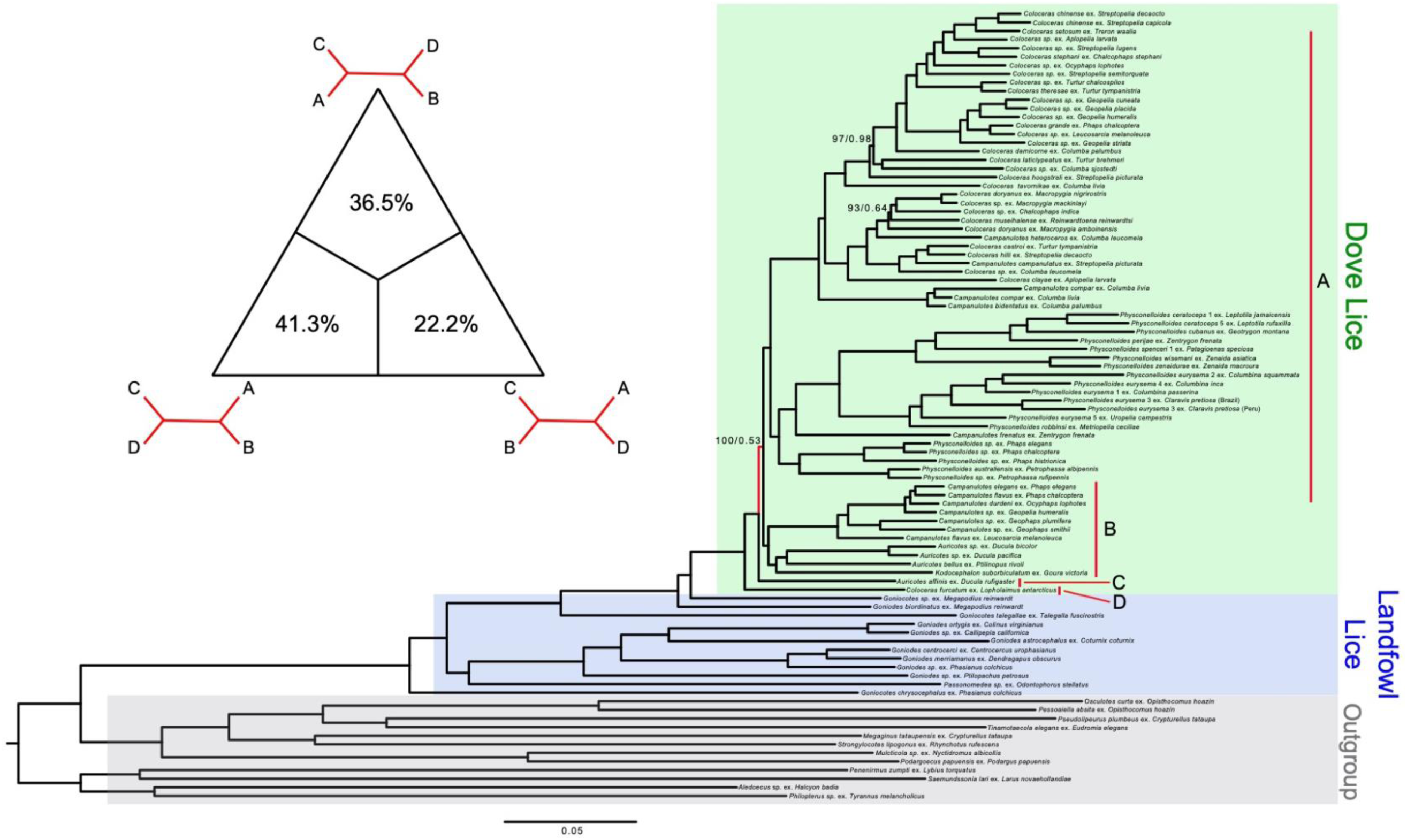
Maximum likelihood phylogeny of dove body lice, landfowl body lice, and outgroup taxa estimated from a concatenated alignment of 2,367 nuclear genes (maximum 60% missing data at each site). Branch support values are shown above branches with less than 100 bootstrap/1.0 local posterior probability. Scale bar indicates branch lengths as nucleotide substitutions per site. The inset figure shows results from a likelihood mapping analysis testing phylogenetic signal at a branch with relatively low support (branch colored red in the phylogeny). The four clades included in the likelihood mapping are indicated on the phylogeny (A-D). Percentages in the triangle indicate the percent of quartets supporting a particular topology

Despite most branches being very highly supported and stable to method of analysis across the phylogeny of body lice and their relatives, a small number of branches did show conflicting phylogenetic signals. *Coloceras museihalense* from *Reinwardtoena reinwardtsi* was recovered as sister to a clade of three other species of *Coloceras* (*C. doryanus* from *Macropygia nigrirostris*, *C.* sp. from *Macropygia mackinlayi*, and *C.* sp. from *Chalcophaps indica*) but with low support (<95% BS and SH-aLRT, <0.9 LPP) in the concatenated and coalescent trees (Figure 1, Supplementary Figure S2). Removing the third codon positions resulted in *C. museihalense* being sister to a clade that included the same three species of *Coloceras* plus *C. doryanus* from *Macropygia amboinensis* (with low support), further suggesting the placement of *C. museihalense* is unstable (Supplementary Figure S8-S12). Another branch with conflicting phylogenetic signals involves *Auricotes affinis* from *Ducula rufigaster*. Concatenated analyses places *A. affinis* as sister to the rest of dove body lice excluding *Coloceras furcatum* with high support (100 BS/100 SH-aLRT). However, this relationship was not supported by most sites (30.3%) or genes (2.18%), and some coalescent analyses recovered *A. affinis* in a clade with *Auricotes, Kodocephalum,* and some species of *Campanulotes* (with low support, <0.8 LPP) (Supplementary Figure S2-S3). Additionally, likelihood mapping indicated 41.3% of quartets supported the relationship recovered from the concatenated analyses, but 58.7% of quartets supported one of the other two arrangements (22.2% for *A. affinis* with clade of *Auricotes, Kodocephalum,* and some *Coloceras*; 36.5% with a clade of *Campanulotes, Coloceras,* and *Physconelloides*) (Figure 1).

Divergence time estimation with MCMCTree and IQ-TREE yielded consistent results (Supplementary Figures S13-S15). MCMCTree generated a phylogeny with an estimated root age of 59.8 MY (52.3-67.9 95% HPD), a stem landfowl louse-dove louse clade age of 58.6 MY (50.9-66.7 MY), and a stem dove body louse age of 26.1 MY (19.2-31.8 MY). Body lice of doves began to diversify (crown body lice) at 22.2 MY (15.3-26.9 MY). The second MCMCTree run was consistent with the first run, with a root age of 60.8 MY (54.1-69.2 MY), stem landfowl louse-dove louse clade age of 59.8 MY (53.0-68.3 MY), stem dove body louse age of 24.5 MY (19.9-29.7 MY), and crown dove body lice at 18.8 MY (15.8-22.5 MY) (Supplementary Figure S14). The tree dated with the LSD2 method in IQ-TREE resulted in a phylogeny with a stem landfowl louse-dove louse clade age of 55.2 MY (52.6-57.2 MY CI), stem dove body louse age of 16.2 MY (14.0-19.7 MY), and crown dove body lice at 13.5 MY (1 2.0-16.2 MY) (Supplementary Figure S15).

### Historical biogeography

Comparisons among several biogeographic models in BioGeoBEARS indicated DIVALIKE+J was the best model for our data (Table S5). The historical biogeographic reconstruction based on the DIVALIKE+J model strongly supported that dove body lice originated in Australasia (Figure 2). There were then subsequent dispersal events from Australia into South America, Eurasia, and Africa, from Eurasia into Africa, from Africa back into Eurasia and into Australasia, and from South America into North America. The best model without the J parameter was the DEC model. Historical biogeographic reconstructions based on DEC also recovered an Australasian origin with high likelihood. The DEC model also recovered multiple dispersal events among different biogeographic regions, but with periods of shared distribution among regions followed by vicariance (Supplementary Figure S16).

**Figure 2.**
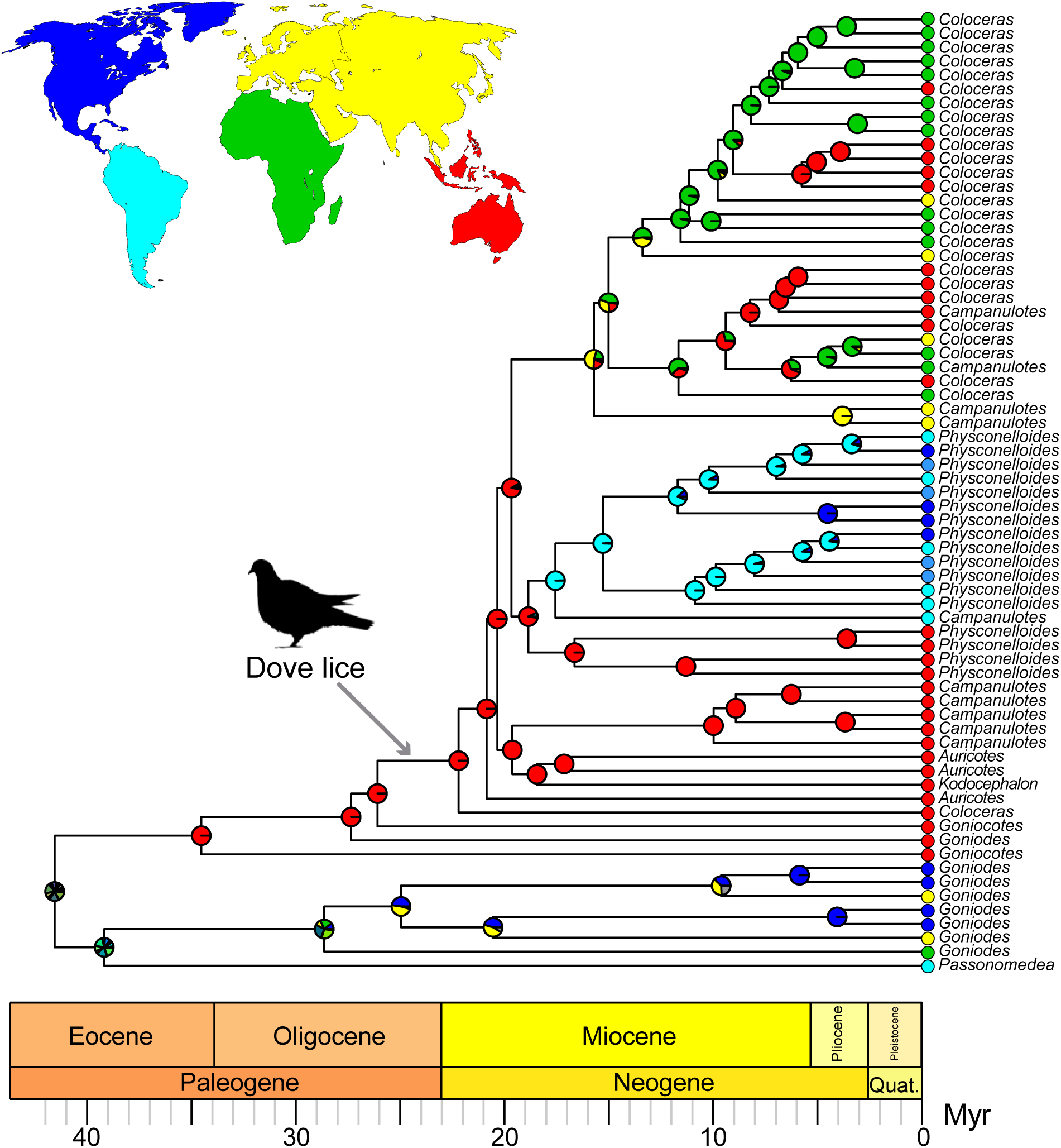
Dated phylogenetic tree of dove body lice and landfowl body lice. Branch lengths are scaled to millions of years. Gray arrow indicates the presumed origin of body lice on doves, as a result of switching from landfowl. Dove silhouette obtained at phylopic.org courtesy of Dori and Nevit Dilmen under a Creative Commons license (https://creativecommons.org/licenses/by-sa/3.0/). Circles at the tips show the current known distributions of that taxon according to one of five biogeographic regions: Africa (green), Australasia (red), Eurasia (yellow), North America (dark blue), South America (light blue). Taxa found in both North and South America are shown with a medium-dark blue. Pie charts at each node indicate the likelihood the ancestor lived in a particular biogeographic region, according to a historical biogeographic reconstruction with BioGeoBEARS under a DIVALIKE+J model.

### Cophylogenetic comparisons

Comparing the trimmed body louse phylogeny (trimmed to OTUs) to the dove phylogeny in Jane generated 28,485 solutions (total cost = 86). The reconciliations included 19-21 cospeciations, 1-3 duplications, 34-36 host switches, 7-10 losses, and 6 failures to diverge (Figure 3, Table S6). Randomizing the tip mappings produce zero solutions with a cost less than the observed cost (*P* < 0.01). Our PACo test also indicated the body louse and dove phylogenies are significantly congruent (m^2^ = 7168.5, p < 0.001). Comparing the body louse tree with the wing louse tree from Boyd et al. (2017) indicated the two louse phylogenies are significantly congruent when trimmed to have overlapping host taxa (m^2^ = 0.49, p < 0.001). Randomizing the wing louse and body louse phylogenies also indicated the observed residual is significantly less than with random phylogenies (p < 0.001) (Supplementary Figure S17-S18).

**Figure 3.**
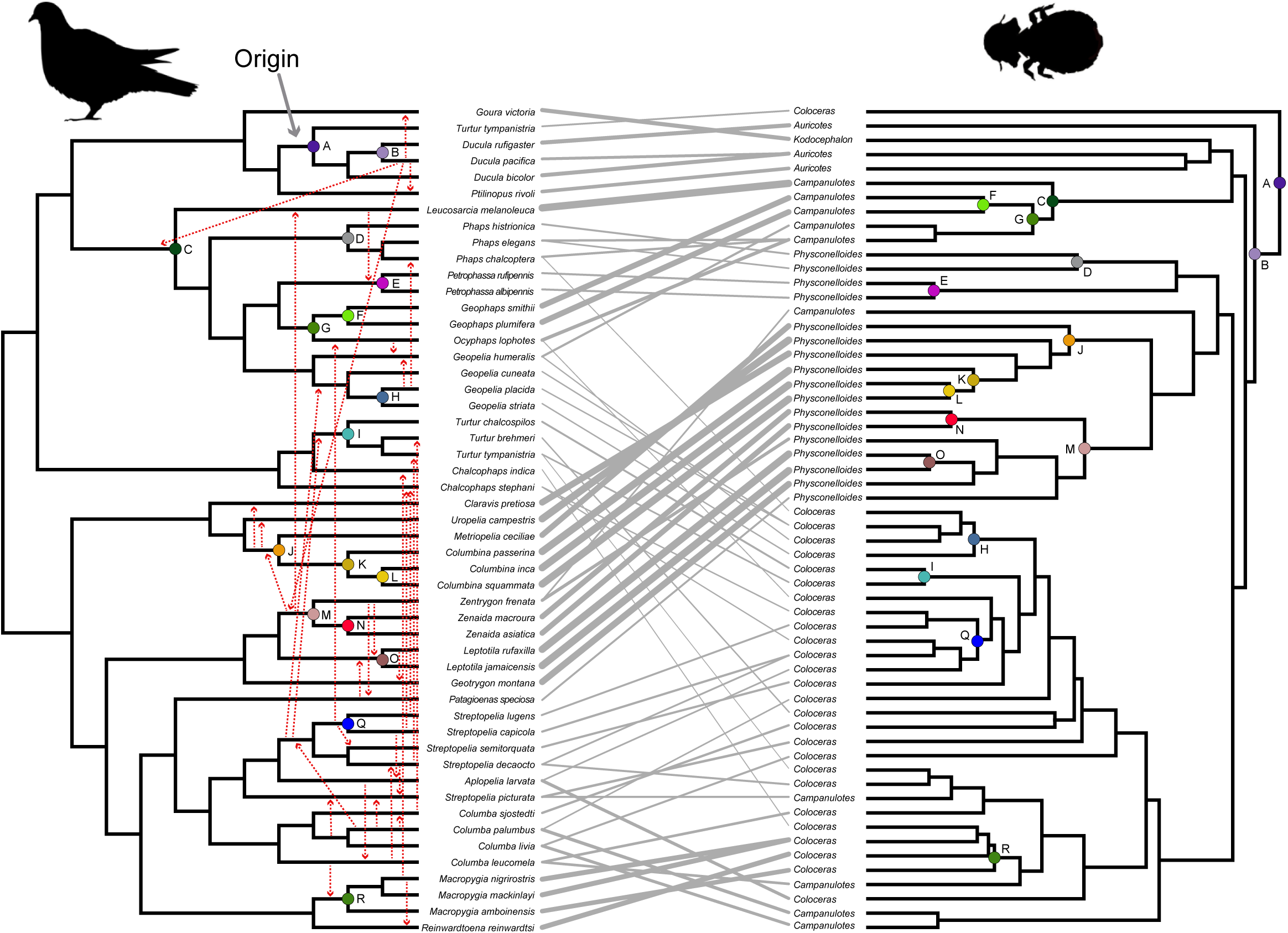
Tanglegram showing the cladograms of doves (left) and their body lice (right). Associated host and parasites are connected by gray lines. The trees are rotated to minimize crossings of the connecting lines. Thickness of the connecting lines indicates the proportional contributions of individual host-parasite associations to the overall congruence between the two trees, as estimated in PACo. Thicker lines indicate a greater contribution to the congruence. Host switches and cospeciations recovered from a reconciliation analysis in Jane are shown as dotted red arrows on the host tree (host switches) and circles (cospeciations). Corresponding cospeciation events are indicated with matching circle colors and letters. The origin of body lice on doves as recovered by the Jane analysis is indicated with a solid gray arrow. Dove silhouette obtained at phylopic.org courtesy of Dori and Nevit Dilmen under a Creative Commons license (https://creativecommons.org/licenses/by-sa/3.0/).

**Figure 4.**
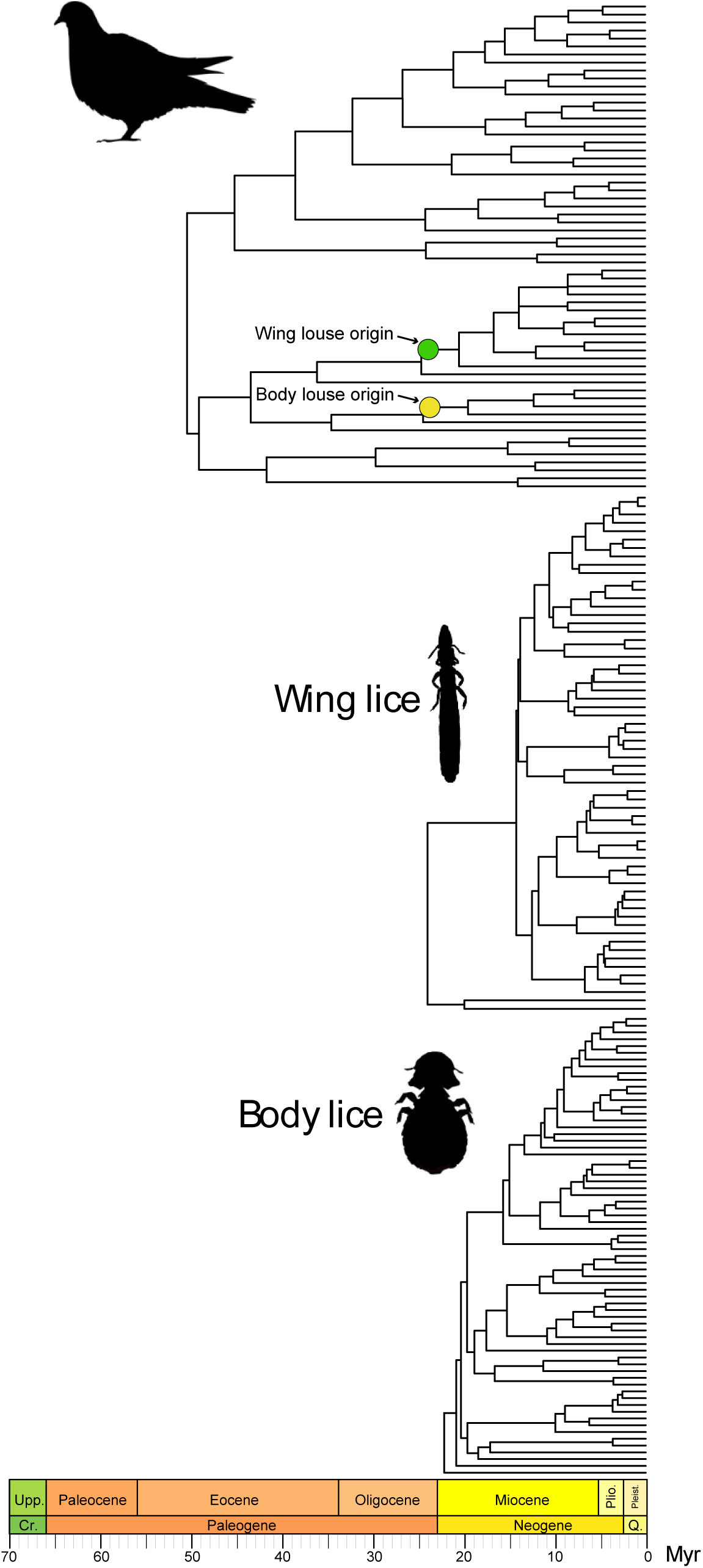
Comparisons of dated phylogenies of doves, dove wing lice, and dove body lice. The dove and wing louse phylogenies are modified from Boyd et al. (2022) and the body louse phylogeny is from the current study. The origins of wing and body lice are shown on the dove phylogeny, based on phylogenetic reconciliation analyses in Jane. Dove silhouette obtained at phylopic.org courtesy of Dori and Nevit Dilmen under a Creative Commons license (https://creativecommons.org/licenses/by-sa/3.0/).

## DISCUSSION

### A single origin of body lice on doves

We found strong support that dove body lice originated once within the lice of landfowl, likely from a host switch from landfowl (Galliformes) to doves (Columbiformes). All our phylogenetic analyses recovered dove body lice as a monophyletic group. On the surface, such a host switch seems unlikely. Galliformes and Columbiformes are not closely related Orders of birds and can have very different natural histories. For instance, our cophylogenetic analysis recovered the ancestor of *Lopholaimus antarcticus* (Topknot pigeon) and *Ducula* (a genus of fruit pigeons) as the original hosts of dove body lice. However, these extant species of doves are arboreal, whereas landfowl are typically ground-dwelling (Gibbs et al. 2001). Members of Galliformes also tend to have larger body sizes than those of Columbiformes, although there is extensive variation in both groups with some overlap. Host body size can be a limiting factor for host suitability in parasitic lice (Bush and Clayton 2006, Villa et al. 2019).

Despite these apparent differences between the two host groups, there are also several lines of evidence that support the feasibility of a switch between these two groups. First, it is possible the ancestor of *L. antarcticus* and *Ducula* was ground-dwelling. There are many species of doves that spend most of their time on the ground, which could create opportunities for exchanging lice with landfowl during times of proximity between the different hosts, such as foraging in mixed groups (Goodwin 1983, Gibbs et al. 2001). Some lice can also be potentially exchanged via the ground itself, e.g., through shared dust baths (Clayton et al. 2004). There is also a size overlap between some doves and landfowl. For example, *L. antarcticus* pigeons can be as large as 600 g and *Megapodius reinwardt*, whose louse *Goniodes biordinatus* is sister to the entire clade of dove body lice, can be as small as 500 g in females (Gibbs et al. 2001, Elliott et al. 2020). Furthermore, comparative studies have shown that body lice are likely less constrained by host body size compared to other types of lice, including wing lice from doves (Johnson et al. 2005). A lack of host size constraints could allow for switches between distantly related hosts. Biogeography also aligns well with a host switching event from Galliformes to Columbiformes. The landfowl hosts of the lice that are sister to the dove louse clade are all distributed in Australiasia, as are those of the earliest diverging dove lice. This biogeographic overlap likely facilitated a host switch between the two groups of hosts. Finally, host switches by lice between distantly related hosts has been documented in several cases. For example, lemur lice in the genus *Trichophilopterus* switched from a bird host (Johnson et al. 2018), and some lice in the *Degeeriella*-complex likely switched between falcons and woodpeckers (Catanach and Johnson 2015).

Our finding of a single origin of dove body lice contradicts the phylogenetic tree of Johnson et al. (2011), who found evidence of a switch from landfowl to doves, but also a switch back to landfowl from doves. A consensus of parsimony trees from Johnson et al. (2011) placed *G. biordinatus* from *M. reinwardt* (Orange-footed Scrubfowl) embedded within the dove body louse phylogeny as sister to a clade of body lice from New World pigeons, Australasian phabine doves, and Australasian fruit doves. A Bayesian phylogeny from Johnson et al. (2011) placed *G. biordinatus* as sister to *Campanulotes frenatus* from *Geotrygon frenata* (White-throated quail-dove). However, these relationships from Johnson et al. (2011) were not well supported (requiring breaking only branches at or below 50% bootstrap and 0.80 Bayesian posterior probability to place *G. biordinatus* outside lice from doves). We also sampled *Goniodes biordinatus*, thus making our taxon sample comparable to the prior Sanger study, but this louse was well outside the clade comprising all the body louse of doves, which were united by a long stem branch with 100% bootstrap and LPP support across all analyses. However, we did recover both *Gonoides* and *Goniocotes* from *Megapodius* as the closest relatives in a clade leading to the body lice of doves, suggesting a close evolutionary relationship between some landfowl lice and dove body lice.

Differences between our phylogenomic tree and the trees from the Sanger data of Johnson et al. (2011) are likely due to gene tree discordance and/or poor phylogenetic signal from individual genes. Many of the backbone branches of the Sanger-based tree were not well supported, indicating a few mitochondrial and nuclear genes are not informative enough for resolving deeper branches in the dove body louse phylogeny. Nevertheless, we do recover dove lice as monophyletic with strong support (99 BS) in our phylogeny based on *cox1* (Supplementary Figure S19), although several of the other relationships differ from our nuclear trees with varying levels of support. This *cox1* data is the full gene sequence (1,575 bp), unlike the previously published Sanger data for this gene, which was only 379 bp long. Thus, even slight increases in the amount of data for this gene can potentially provide increased resolution.

Variation in topologies is also likely driven by variation in the evolutionary histories at loci across a genome or between mitochondrial and nuclear loci. Our analyses of thousands of nuclear genes support this variation. Even though we had high support for most of the branches throughout our phylogenetic trees, some branches had conflicting support depending on the analytical approach. Most notably, a branch splitting *Auricotes affinis* and *Coloceras furcatum* from the rest of dove body lice received high bootstrap and SH-aLRT support but low support from coalescent analyses. Coalescent analyses without third codon positions also recovered an alternative topology, though with low support. Because of this discrepancy between concatenated and coalescent approaches, we expected considerable variation in gene trees at this branch. Consistent with this expectation, we found that only 2.18% of gene trees and 30.3% of sites supported the topology recovered supported by the concatenated analysis, suggesting considerable gene tree-species tree conflict. Conflict between gene trees and the species tree can result from different biological processes, including ancestral hybridization and Incomplete Lineage Sorting (ILS) (Maddison 1997). A previous study has found evidence for introgression due to hybridization and backcrossing in dove lice (Doña et al. 2020). The branch splitting *A. affinis* and *Coloceras furcatum* from other lice is very short, and thus much more likely to be affected by ILS. If the ancestral effective population size was large, this would greatly increase the probability of genes not sorting according to species during a period of relatively rapid diversification.

### Dove body lice originated in Australasia and dispersed across the world with their hosts

Body lice from doves likely switched from landfowl, possibly megapodes, in Australasia. Our historical biogeographic reconstruction recovered Australasia as the ancestral range of dove body lice around 23-26 MYA (>99% likelihood). The likely original hosts, the ancestors of *L. antarcticus* and *Ducula* sp., were also likely present in Australasia at this time. Both *L. antarcticus* and *Ducula* sp. have extant ranges in Australasia and last shared a common ancestor ∼20 MYA (Boyd et al. 2022).

Soon after originating on doves, body lice started spreading across the globe. A burst of rapid divergence ∼20 MYA, indicated by several short branches in the phylogeny (Figure 2), suggests the range expansion began relatively quickly. Rapid expansion is consistent with the biogeographic and evolutionary history of their dove hosts. Doves likely originated in Australasia/South America before the Gondwanan breakup. Around 20 MYA, doves began to expand across the globe, including dispersal out of Australasia. Some doves also subsequently dispersed back into Australasia from Eurasia and the western hemisphere (Boyd et al. 2022). The biogeographic history of dove lice seems to reflect this pattern of back-dispersal, with several instances of dispersal back into Australasia in the lice.

Dispersal out of Australasia in the early Miocene has been found in other groups of organisms. For example, the Corvides clade of passerine birds dispersed out of Australasia multiple times in the early Miocene (Oliveros et al. 2019). Similar biogeographic patterns have been found in a wide range of taxa, including a subfamily of freshwater snails (Miratestinae) (Gauffre-Autelin et al. 2021) and plants dispersed by animals (Grudinksi et al. 2014). There are several plausible explanations for why doves and other organisms dispersed widely in the early Miocene. Dispersal could have been driven by major geological changes occurring in the Indo-Australasian peninsula during this time period (de Bruyn et al. 2014). There were also major climatic changes during the Miocene, including drastic cooling and drying across many continents in the late Miocene (Herbert et al. 2016). Both geologic and climatic changes could have led to more suitable habitat for doves, such as expanded C4 grasslands in the late Miocene (Osborne 2007). Although many doves are arboreal frugivores, several lineages are primarily terrestrial seed-eaters, including independent lineages occupying African, New World, and Australian grasslands, for which expansion of grasslands may have facilitated their diversification. Regardless of why doves began dispersing around 20 MYA, our results highlight that this dispersal also drove global dispersal of dove body lice, which could have facilitated subsequent host switches among hosts that had been previously separated by time and geography (Brooks et al. 2019), and this also seems to be the case for the wing lice of doves (Boyd et al. 2022).

Future work could help tease apart the fine-scale dispersal patterns of doves and their lice by including a larger taxon sample focused on Australasia and Eurasia. Our current study shows strong support for the reconstruction of broad biogeographic patterns, but we are unable to reconstruct some dispersal events. For example, we do not have a dense sampling of lice from Eurasia (relative to Australasia and Africa). Including more samples from Eurasia could help connect dispersal events form Australiasia to the New World and Africa back into Australasia. Both dispersal events may have had had stepping-stones in Eurasia, which could be more evident with a denser sampling in the Eurasian biogeographic region, particularly from East Asia and the Indian subcontinent.

### Dove body and wing lice have similar biogeographic histories

The diversification and biogeographic histories of dove body lice have many consistencies with the evolutionary history of dove wing lice (Figure 5). Both body and wing lice likely originated on Australasian doves ∼25 MYA and soon thereafter dispersed across the world with their hosts. The phylogenies of wing and body lice were congruent with one another, suggesting long-term evolutionary similarities between the two groups of lice, possibly driven by underlying patterns of codivergence with a shared host group. This is supported by the fact that body and wing lice both showed evidence of codivergence with their hosts. Both groups also experience host-switches to other hosts after dispersing among biogeographic regions. This consistency is surprising given that the two groups of lice are not closely related and have different dispersal abilities. Previous work has shown that wing lice, which are better able to disperse by hitchhiking on winged hippoboscid flies, tend to switch hosts more frequently than body lice (Clayton and Johnson 2003). As a result, wing lice typically have less phylogenetic congruence with their hosts (Clayton and Johnson 2003, but see Sweet et al. 2017), less population structure, and larger effective population sizes compared to body lice (Sweet and Johnson 2018). Although these prior results likely accurately reflect coevolutionary relationships, our results suggest that host-related factors can be the most important factor for shaping the diversification and biogeographic history of their parasites at deep evolutionary timescales. Similarities in the cophylogenetic history of body and wing lice with doves could also be a result of more general evolutionary process, such as competition for available resources.

Another line of comparison is the relative contribution of host-switching versus cospeciation to parasite speciation events over time. Boyd et al. (2022) found a decrease in the proportion of speciation events in dove wing lice attributed to host-switching versus cospeciation over time. This decrease was gradual from about 85% of nodes among early diverging lineages to 50% of nodes attributed to host-switching at around 5 MYA. As doves dispersed into new geographic areas, there could have been more ecological opportunities for wing lice to move to new hosts already in the region (or vice versa), facilitating host switching. Over time the niches would be filled by similar species of parasites, which could make it challenging for a louse to establish populations on new species of hosts, thus deterring host switching and promoting codivergence over time (Agostal et al. 2010, Brooks et al. 2019, Boyd et al. 2022). Unlike dove wing lice, however, our results suggest that the relative proportion of speciation events attributable to host-switching in the body lice of doves is relatively constant at around 50% to 60% over all time periods. Thus, both host-switching and cospeciation occurred with similar frequencies regardless of the ecological context (Supplementary Figure S18). Because of their lower dispersal capabilities, it may be that dove body lice are more likely to cospeciate with their hosts even if there are open niches of dove lineages as yet uncolonized by body lice. Over time, body lice may fill these open niches, but less rapidly than wing lice because of their lower dispersal ability. Thus, while host-switching dominates early divergence events in wing lice, it declines to a similar proportion to that in body lice over time.

### Implications for generic level taxonomy of dove body lice

Our phylogenetic analyses indicate the four genera (following Price et al. 2003) of dove body lice sampled by more than one representative are not monophyletic. Both our nuclear and *cox1* trees recovered each of these four genera (*Coloceras*, *Campanulotes*, *Physconelloides*, and *Auricotes*) as paraphyletic. This finding is consistent with prior studies (Johnson et al. 2001, 2011), which also failed to recover monophyletic genera. However, some authors (Tendeiro 1969, 1971) have split the genus *Campanulotes* into additional genera, including *Saussurites* for lice from Australian phabines and New World doves, and *Nitzschielloides* for *Campanulotes campanulatus*. However, even in this scheme *Saussurites* would be paraphyletic, with *Campanulotes* (*Saussurites*) *frenatus* being separated in our tree from other *Campanulotes* (which also renders *Physconelloides* paraphyletic). The genus *Campanulotes* is generally recognized by its small body size, and thus reduction in characters might be leading to superficial morphological similarity. The same is true of the genera *Goniodes* (large-bodied) and *Goniocotes* (small-bodied) from landfowl (Galliformes), that even with our limited we find as highly paraphlytic. Nevertheless, our results further strengthen the conclusions of previous work suggesting the genus-level taxonomy of Goniodidae needs to be reevaluated. The well resolved and highly supported phylogenomic tree of this present study provides an important new framework under which morphological evidence for generic limits can be reevaluated in this group.

## ACKNOWLEDGMENTS

We are grateful for the following individuals for providing samples of lice: John Bates, Sarah Bush, Terry Chesser, Dale Clayton, Rob Faucett, A.W. Krater, Jael Malenke, Ian Mason, Kevin McCracken, Vitor de Q. Piacentini, Vince Smith, Michel Valim, Dave Willard, and Rob Wilson. We also thank Alvaro Hernandez and Chris Wright at the Roy J. Carver Biotechnology Center at the University of Illinois for their efforts to generate genome sequence data from our samples. We thank Kimberly K. O. Walden for assistance with submission of raw reads to NCBI. We also thank Paige Brewer for assistance with phylogenetic analyses, Bret Boyd for discussions about biogeography and dispersal, and the Arkansas High Performance Computing Center at the University of Arkansas for providing computational resources.

## DATA AVAILABILITY

Raw genome sequence data is available at the NCBI SRA database. Data files and supplementary material can be found in the Dryad data repository: (accession # pending).

## LITERATURE CITED

Agosta S.J., Janz N., Brooks D.R. 2010. How specialists can be generalists: resolving the “parasite paradox” and implications for emerging infectious disease. Zoologia (Curitiba). 27:151–162. 10.1590/S1984-46702010000200001.

Allen J.M., LaFrance R., Folk R.A., Johnson K.P., Guralnick R.P. 2018. aTRAM 2.0: An improved, flexible locus assembler for NGS data. Evolutionary Bioinformatics. 14:117693431877454. 10.1177/1176934318774546

Balbuena J.A., Míguez-Lozano R., Blasco-Costa I. 2013. PACo: A novel procrustes application to cophylogenetic analysis. PLoS ONE. 8. 10.1371/journal.pone.0061048

Barrett L.G., Thrall P.H., Burdon J.J., Linde C.C. 2008. Life history determines genetic structure and evolutionary potential of host--parasite interactions. Trends in Ecology & Evolution. 23:678–685. 10.1016/j.tree.2008.06.017

Benovics M., Desdevises Y., Šanda R., Vukić J., Šimková A. 2020. Cophylogenetic relationships between *Dactylogyrus* (Monogenea) ectoparasites and endemic cyprinoids of the north-eastern European peri-Mediterranean region. Journal of Zoological Systematics and Evolutionary Research. 58:1–21. 10.1111/jzs.12341

Blasco-Costa I., Hayward A., Poulin R., Balbuena J.A. 2021. Next-generation cophylogeny: unravelling eco-evolutionary processes. Trends in Ecology and Evolution. 36:907–918. 10.1016/j.tree.2021.06.006

Bordes F., Morand S. 2009. Parasite diversity: an overlooked metric of parasite pressures? Oikos. 118:801–806. 10.1111/j.1600-0706.2008.17169.x

Borowiec M.L. 2016. AMAS: a fast tool for alignment manipulation and computing of summary statistics. PeerJ. 4:e1660. 10.7717/peerj.1660

Boyd B.M., Allen J.M., Nguyen N., Sweet A.D., Warnow T., Shapiro M.D., Villa S.M., Bush S.E., Clayton D.H., Johnson K.P. 2017. Phylogenomics using target-restricted assembly resolves intra-generic relationships of parasitic lice (Phthiraptera: *Columbicola*). Systematic Biology. 66:896–911. 10.1093/sysbio/syx027

Boyd B.M., Nguyen N.P., Allen J.M., Waterhouse R.M., Vo K.B., Sweet A.D., Clayton D.H., Bush S.E., Shapiro M.D., Johnson K.P. 2022. Long-distance dispersal of pigeons and doves generated new ecological opportunities for host-switching and adaptive radiation by their parasites. Proceedings of the Royal Society B: Biological Sciences. 289. 10.1098/rspb.2022.0042

Brooks D.R., Hoberg E.P., Boeger W.A. 2019. The Stockholm Paradigm: climate change and emerging disease. The Stockholm Paradigm. University of Chicago Press.

de Bruyn M., Stelbrink B., Morley R.J., Hall R., Carvalho G.R., Cannon C.H., van den Bergh G., Meijaard E., Metcalfe I., Boitani L., Maiorano L., Shoup R., von Rintelen T. 2014. Borneo and Indochina are major evolutionary hotspots for Southeast Asian biodiversity. Systematic Biology. 63:879–901. 10.1093/sysbio/syu047

Bush S.E., Clayton D.H. 2006. The role of body size in host specificity: Reciprocal transfer experiments with feather lice. Evolution. 60:2158–2167. 10.1111/j.0014-3820.2006.tb01853.x

Bush S.E., Malenke J.R. 2008. Host defence mediates interspecific competition in ectoparasites. Journal of Animal Ecology. 77:558–564. 10.1111/j.1365-2656.2007.01353.x

Cailliez F. 1983. The analytical solution of the additive constant problem. Psychometa. 48:305– 308.

Capella-Gutiérrez S., Silla-Martínez J.M., Gabaldón T. 2009. trimAl: a tool for automated alignment trimming in large-scale phylogenetic analyses. Bioinformatics. 25:1972–1973. 10.1093/bioinformatics/btp348

Catanach T.A., Johnson K.P. 2015. Independent origins of the feather lice (Insecta: Degeeriella) of raptors. Biological Journal of the Linnean Society. 114:837–847. 10.1111/bij.12453

Chen S., Zhou Y., Chen Y., Gu J. 2018. fastp: an ultra-fast all-in-one FASTQ preprocessor. Bioinformatics. 34:i884–i890. 10.1093/bioinformatics/bty560

Cibois A., Thibault J.C., Bonillo C., Filardi C.E., Watling D., Pasquet E. 2014. Phylogeny and biogeography of the fruit doves (Aves: Columbidae). Molecular Phylogenetics and Evolution. 70:442–453. 10.1016/j.ympev.2013.08.019

Clay T. 1949. Some Problems in the Evolution of a Group of Ectoparasites. Evolution. 3:279– 299.

Clayton D.H. 1991. Coevolution of avian grooming and ectoparasite avoidance. In: Loye J.E., Zuk M., editors. Bird-parasite interactions: ecology, evolution, and behavior. Oxford: Oxford University Press. p. 258–289.

Clayton D.H., Bush S.E., Goates B.M., Johnson K.P. 2003. Host defense reinforces host–parasite cospeciation. PNAS. 100:15694–15699. 10.1073/pnas.2533751100

Clayton D.H., Bush S.E., Johnson K.P. 2004. Ecology of congruence: Past meets present. Systematic Biology. 53:165–173. 10.1080/10635150490265102

Clayton D.H., Bush S.E., Johnson K.P. 2016. Coevolution of life on hosts: integrating ecology and history. Chicago: The University of Chicago Press.

Clayton D.H., Drown D.M. 2001. Critical evaluation of five methods for quantifying chewing lice (Insecta: Phthiraptera). Journal of Parasitology. 87:1291–1300. 10.2307/3285290

Clayton D.H., Johnson K.P. 2003. Linking coevolutionary history to ecological process: doves and lice. Evolution. 57:2335–41. 10.1111/j.0014-3820.2003.tb00245.x

Clayton D.H., Walther B.A.. 1997. Collection and quantification of arthropod parasites of birds. Host-parasite evolution: general principles and avian models. In: Host-parasite evolution: general principles and avian models. Clayton D.H. and Moore J., editors. Oxford University Press, Oxford. p. 419–440.

Conow C., Fielder D., Ovadia Y., Libeskind-Hadas R. 2010. Jane: a new tool for the cophylogeny reconstruction problem. Algorithms for Molecular Biology. 5:16.. 10.1186/1748-7188-5-16

Cruaud A., Ronsted N., Chantarasuwan B., Chou L.S., Clement W.L., Couloux A., Cousins B., Genson G., Harrison R.D., Hanson P.E., Hossaert-Mckey M., Jabbour-Zahab R., Jousselin E., Kerdelhué C., Kjellberg F., Lopez-Vaamonde C., Peebles J., Peng Y.Q., Santinelo Pereira R.A., Schramm T., Ubaidillah R., Van Noort S., Weiblen G.D., Yang D.R., Yodpinyanee A., Libeskind-Hadas R., Cook J.M., Rasplus J.Y., Savolainen V. 2012. An extreme case of plant-insect codiversification: Figs and fig-pollinating wasps. Systematic Biology. 61:1029–1047. 10.1093/sysbio/sys068

De Vienne D.M., Giraud T., Shykoff J.A. 2007. When can host shifts produce congruent host and parasite phylogenies? A simulation approach. Journal of Evolutionary Biology. 20:1428–1438. 10.1111/j.1420-9101.2007.01340.x

Diblasi E., Johnson K.P., Stringham S.A., Hansen A.N., Beach A.B., Clayton D.H., Bush S.E. 2018. Phoretic dispersal influences parasite population genetic structure. Molecular Ecology. 27:2770–2779. 10.1111/mec.14719

Dismukes W., Braga M.P., Hembry D.H., Heath T.A., Landis M.J. 2022. Cophylogenetic methods to untangle the evolutionary history of ecological interactions. Annual Review of Ecology, Evolution, and Systematics. 53:275–298. 10.1146/annurev-ecolsys-102320-112823

Doña J., Sweet A.D., Johnson K.P. 2020. Comparing rates of introgression in parasitic feather lice with differing dispersal capabilities. Communications Biology. 3:610. 10.1038/s42003-020-01345-x

Eichler W.D. 1948. Some rules in ectoparasitism. The Annals and Magazine of Natural History. 1:588–598.

Elliott, A., Kirwan, G.M., Christie, D.A. 2020. Orange-footed Megapode (*Megapodius reinward*t), version 1.0. In Birds of the World (J. del Hoyo, A. Elliott, J. Sargatal, D. A. Christie, and E. de Juana, Editors). Cornell Lab of Ornithology, Ithaca, NY, USA. 10.2173/bow.orfscr1.01

Ellis V.A., Collins M.D., Medeiros M.C.I., Sari E.H.R., Coffey E.D., Dickerson R.C., Lugarini C., Stratford J.A., Henry D.R., Merrill L., Matthews A.E., Hanson A.A., Roberts J.R., Joyce M., Kunkel M.R., Rickelfs R.E.. 2015. Local host specialization, host-switching, and dispersal shape the regional distributions of avian haemosporidian parasites. PNAS. 112:11294–11299. 10.1073/pnas.1515309112

Fahrenholz H. 1913. Ectoparasiten und abstammungslehre. Zoologischer Anzeiger. 41:371–374.

Gauffre-Autelin P., Stelbrink B., von Rintelen T., Albrecht C. 2021. Miocene geologic dynamics of the Australian Sahul Shelf determined the biogeographic patterns of freshwater planorbid snails (Miratestinae) in the Indo-Australian Archipelago. Molecular Phylogenetics and Evolution. 155:107004. 10.1016/j.ympev.2020.107004

Gibbs D.E., Cox E., Cox J. 2001. Pigeons and Doves: A guide to the pigeons and doves of the world. Sussex: Pica Press.

Goodwin D. 1983. Pigeons and Doves of the World. Ithaca, New York: Cornell University Press.

Grudinski M., Wanntorp L., Pannell C.M., Muellner-Riehl A.N. 2014. West to east dispersal in a widespread animal-dispersed woody angiosperm genus (*Aglaia*, Meliaceae) across the Indo-Australian Archipelago. Journal of Biogeography. 41:1149–1159. 10.1111/jbi.12280

Guindon S., Dufayard J.-F., Lefort V., Anisimova M., Hordijk W., Gascuel O. 2010. New algorithms and methods to estimate maximum-likelihood phylogenies: Assessing the performance of PhyML 3.0. Systematic Biology. 59:307–321. 10.1093/sysbio/syq010

Harbison C.W., Jacobsen M.V., Clayton D.H. 2009. A hitchhiker’s guide to parasite transmission: The phoretic behaviour of feather lice. International Journal for Parasitology. 39:569–575. 10.1016/j.ijpara.2008.09.014

Harmon L.J., Andreazzi C.S., Débarre F., Drury J., Goldberg E.E., Martins A.B., Melián C.J., Narwani A., Nuismer S.L., Pennell M.W., Rudman S.M., Seehausen O., Silvestro D., Weber M., Matthews B. 2019. Detecting the macroevolutionary signal of species interactions. Journal of Evolutionary Biology. 32:769–782. 10.1111/jeb.13477

Hay E.M., Poulin R., Jorge F. 2020. Macroevolutionary dynamics of parasite diversification: A reality check. Journal of Evolutionary Biology. 33: 1758–1769. 10.1111/jeb.13714

Hembry D.H., Weber M.G. 2020. Ecological interactions and macroevolution: a new field with old roots. Annual Review of Ecology, Evolution, and Systematics. 51:215–243. 10.1146/annurev-ecolsys-011720-121505

Herbert T.D., Lawrence K.T., Tzanova A., Peterson L.C., Caballero-Gill R., Kelly C.S. 2016. Late Miocene global cooling and the rise of modern ecosystems. Nature Geoscience. 9:843–847. 10.1038/ngeo2813

Ho S.Y.W., Duchêne S. 2014. Molecular-clock methods for estimating evolutionary rates and timescales. Molecular Ecology. 23:5947–5965. 10.1111/mec.12953

Hoang D.T., Chernomor O., von Haeseler A., Minh B.Q., Vinh L.S. 2018. UFBoot2: Improving the ultrafast bootstrap approximation. Molecular Biology and Evolution. 35:518–522. 10.1093/molbev/msx281

Hoberg E.P., Brooks D.R. 2008. A macroevolutionary mosaic: episodic host-switching, geographical colonization and diversification in complex host-parasite systems. Journal of Biogeography. 35:1533–1550. 10.1111/j.1365-2699.2008.01951.x

Hughes J., Kennedy M., Johnson K.P., Palma R.L., Page R.D.M. 2007. Multiple cophylogenetic analyses reveal frequent cospeciation between pelecaniform birds and *Pectinopygus* lice. Systematic Biology. 56:232–51. 10.1080/10635150701311370

Hutchinson M.C., Cagua E.F., Balbuena J.A., Stouffer D.B., Poisot T. 2017. paco: implementing Procrustean Approach to Cophylogeny in R. Methods in Ecology and Evolution. 8:932– 940. 10.1111/2041-210X.12736

Jezkova T., Wiens J.J. 2017. What explains patterns of diversification and richness among animal phyla? The American Naturalist. 189:201–212. 10.1086/690194

Johnson, K.P., Adams R.J., Clayton D.H. 2001. Molecular systematics of Goniodidae (Insecta: Phthiraptera). Journal of Parasitology 87: 862–869. 10.1645/0022-3395(2001)087[0862:MSOGIP]2.0.CO;2.

Johnson K.P., Bush S.E., Clayton D.H. 2005. Correlated evolution of host and parasite body size: tests of Harrison’s Rule using birds and lice. Evolution. 59:1744–1753. 10.1111/j.0014-3820.2005.tb01823.x

Johnson K.P., Dietrich C.H., Friedrich F., Beutel R.G., Wipfler B., Peters R.S., Allen J.M., Petersen M., Donath A., Walden K.K.O., Kozlov A.M., Podsiadlowski L., Mayer C., Meusemann K., Vasilikopoulos A., Waterhouse R.M., Cameron S.L., Weirauch C., Swanson D.R., Percy D.M., Hardy N.B., Terry I., Liu S., Zhou X., Misof B., Robertson H.M., Yoshizawa K. 2018. Phylogenomics and the evolution of hemipteroid insects. PNAS. 115:12775–12780. 10.1073/pnas.1815820115

Johnson K.P., Clayton D.H., Dumbacher J.P., Fleischer R.C. 2010. The flight of the Passenger Pigeon: phylogenetics and biogeographic history of an extinct species. Molecular Phylogenetics and Evolution. 57:455–8. 10.1016/j.ympev.2010.05.010

Johnson K.P., Matthee C., Doña J. 2022. Phylogenomics reveals the origin of mammal lice out of Afrotheria. Nature Ecology and Evolution. 6:1205–1210. 10.1038/s41559-022-01803-1

Johnson K.P., Nguyen N.-P., Sweet A.D., Boyd B.M., Warnow T., Allen J.M. 2018. Simultaneous radiation of bird and mammal lice following the K-Pg boundary. Biology Letters. 14:20180141. 10.1098/rsbl.2018.0141

Johnson K.P., Clayton D.H. 2000. A molecular phylogeny of the dove genus *Zenaida*: Mitochondrial and nuclear DNA sequences. The Condor. 102:864–870. 10.1093/condor/102.4.864

Johnson K.P., Weckstein J.D. 2011. The Central American land bridge as an engine of diversification in New World doves. Journal of Biogeography. 38:1069–1076. 10.1111/j.1365-2699.2011.02501.x

Johnson K.P., Weckstein J.D., Virrueta Herrera S., Doña J. 2021. The interplay between host biogeography and phylogeny in structuring diversification of the feather louse genus *Penenirmus*. Molecular Phylogenetics and Evolution. 165:107297. 10.1016/j.ympev.2021.107297

Justine J.-L., Briand M.J., Bray R.A. 2012. A quick and simple method, usable in the field, for collecting parasites in suitable condition for both morphological and molecular studies. Parasitology Research. 111:341–351. 10.1007/s00436-012-2845-6

Kalyaanamoorthy S., Minh B.Q., Wong T.K.F., von Haeseler A., Jermiin L.S. 2017. ModelFinder: fast model selection for accurate phylogenetic estimates. Nature Methods 2017 14:6. 14:587–589. 10.1038/nmeth.4285

Katoh K., Misawa K., Kuma K., Miyata T. 2002. MAFFT: a novel method for rapid multiple sequence alignment based on fast Fourier transform. Nucleic acids research. 30:3059– 3066. 10.1093/nar/gkf436

Katoh K., Standley D.M. 2013. MAFFT Multiple Sequence Alignment Software Version 7: Improvements in performance and usability. Molecular Biology and Evolution. 30:772– 780. 10.1093/molbev/mst010

Koch C.F. 1978. Bias in the published fossil record. Paleobiology. 4:367–372.

Lafferty K.D., Dobson A.P., Kuris A.M. 2006. Parasites dominate food web links. PNAS. 103:11211–11216. 10.1073/pnas.0604755103

Maddison W.P. 1997. Gene trees in species trees. Systematic Biology. 46:523–536. 10.1093/sysbio/46.3.523

Marshall A.G. 1981. The ecology of ectoparasitic insects. Academic Press. Cambridge, MA.

Michelet L., Dauga C. 2012. Molecular evidence of host influences on the evolution and spread of human tapeworms. Biological Reviews. 87:731–741. 10.1111/j.1469-185X.2012.00217.x

Minh B.Q., Hahn M.W., Lanfear R. 2020a. New methods to calculate concordance factors for phylogenomic datasets. Molecular Biology and Evolution. 37:2727–2733. 10.1093/molbev/msaa106

Minh B.Q., Schmidt H.A., Chernomor O., Schrempf D., Woodhams M.D., von Haeseler A., Lanfear R. 2020b. IQ-TREE 2: New models and efficient methods for phylogenetic inference in the genomic era. Molecular Biology and Evolution. 37:1530–1534. 10.1093/molbev/msaa015

Mo Y.K., Lanfear R., Hahn M.W., Minh B.Q. 2023. Updated site concordance factors minimize effects of homoplasy and taxon sampling. Bioinformatics. 39:btac741. 10.1093/bioinformatics/btac741

Morand S. 2015. (macro-) Evolutionary ecology of parasite diversity: From determinants of parasite species richness to host diversification. International Journal for Parasitology: Parasites and Wildlife. 4:80–87. 10.1016/j.ijppaw.2015.01.001

de Moya R.S., Yoshizawa K., Walden K.K.O., Sweet A.D., Dietrich C.H., Kevin P J. 2021. Phylogenomics of parasitic and nonparasitic lice (Insecta: Psocodea): Combining sequence data and exploring compositional bias solutions in next generation data sets. Systematic Biology. 70:719–738. 10.1093/sysbio/syaa075

Nadler S.A., De Leon G.P.P. 2011. Integrating molecular and morphological approaches for characterizing parasite cryptic species: implications for parasitology. Parasitology. 138:1688–1709. 10.1017/S003118201000168X

Nelson B.C., Murray M.D. 1971. The distribution of Mallophaga on the domestic pigeon (*Columba livia*). International Journal for Parasitology. 1:21–29. 10.1016/0020-7519(71)90042-7

Nowak J., Sweet A., Weckstein J., Johnson K. 2019. A molecular phylogenetic analysis of the genera of fruit doves and allies using dense taxonomic sampling. Illinois Natural History Survey Bulletin. 40(2019).

Oliveros C.H., Field D.J., Ksepka D.T., Barker F.K., Aleixo A., Andersen M.J., Alström P., Benz B.W., Braun E.L., Braun M.J., Bravo G.A., Brumfield R.T., Chesser R.T., Claramunt S., Cracraft J., Cuervo A.M., Derryberry E.P., Glenn T.C., Harvey M.G., Hosner P.A., Joseph L., Kimball R.T., Mack A.L., Miskelly C.M., Peterson A.T., Robbins M.B., Sheldon F.H., Silveira L.F., Smith B.T., White N.D., Moyle R.G., Faircloth B.C. 2019. Earth history and the passerine superradiation. PNAS. 116:7916– 7925. 10.1073/pnas.1813206116

Osborne C.P. 2008. Atmosphere, ecology and evolution: what drove the Miocene expansion of C4 grasslands? Journal of Ecology. 96:35–45. 10.1111/j.1365-2745.2007.01323.x

Paradis E., Schliep K. 2019. ape 5.0: an environment for modern phylogenetics and evolutionary analyses in R. Bioinformatics. 35:526–528. 10.1093/bioinformatics/bty633

Park E., Jorge F., Poulin R. 2020. Shared geographic histories and dispersal contribute to congruent phylogenies between amphipods and their microsporidian parasites at regional and global scales. Molecular Ecology. 29:3330–3345. 10.1111/mec.15562

Paterson A.M., Banks J. 2001. Analytical approaches to measuring cospeciation of host and parasites: through a looking glass, darkly. International Journal for Parasitology. 31:1012–1022. 10.1016/S0020-7519(01)00199-0

Paterson A.M., Wallis G.P., Wallis L.J., Gray R.D. 2000. Seabird and louse coevolution: complex histories revealed by 12S rRNA sequences and reconciliation analyses. Systematic Biology. 49:383–99. 10.1080/10635159950127303

Penczykowski R.M., Laine A.-L., Koskella B. 2016. Understanding the ecology and evolution of host--parasite interactions across scales. Evolutionary Applications. 9:37–52. 10.1111/eva.12294

Pereira S.L., Johnson K.P., Clayton D.H., Baker A.J. 2007. Mitochondrial and nuclear DNA sequences support a Cretaceous origin of Columbiformes and a dispersal-driven radiation in the Paleocene. Systematic Biology. 56:656–672. 10.1080/10635150701549672

Poulin R., Morand S. 2000. The diversity of parasites. The Quarterly Review of Biology. 75:277–293. 10.1086/393500

Price R.D., Hellenthal R.A., Palma R.L., Johnson K.P., Clayton D.H. 2003. The chewing lice: world checklist and biological overview. Champaign, IL: Illinois Natural History Survey.

Rambaut A., Drummond A.J., Xie D., Baele G., Suchard M.A. 2018. Posterior summarization in Bayesian phylogenetics using Tracer 1.7. Systematic Biology. 67:901–904. 10.1093/sysbio/syy032

Ree R.H., Sanmartín I. 2018. Conceptual and statistical problems with the DEC+J model of founder-event speciation and its comparison with DEC via model selection. Journal of Biogeography. 45:741–749. 10.1111/jbi.13173

Reis M. dos, Yang Z. 2011. Approximate likelihood calculation on a phylogeny for Bayesian estimation of divergence times. Molecular Biology and Evolution. 28:2161–2172. 10.1093/molbev/msr045

Revell L.J. 2012. phytools: an R package for phylogenetic comparative biology (and other things). Methods in Ecology and Evolution. 3:217–223. 10.1111/j.2041-210X.2011.00169.x

Ricklefs R.E. 2010. Evolutionary diversification, coevolution between populations and their antagonists, and the filling of niche space. PNAS. 107:1265–1272. 10.1073/pnas.0913626107

Robinson D.F., Foulds L.R. 1981. Comparison of phylogenetic trees. Mathematical Biosciences. 53:131–147. 10.1016/0025-5564(81)90043-2

Sayyari E., Mirarab S. 2016. Fast coalescent-based computation of local branch support from quartet frequencies. Molecular biology and evolution. 33:1654–1668. 10.1093/molbev/msw079

Schliep K.P. 2011. phangorn: phylogenetic analysis in R. Bioinformatics. 27:592–593. 10.1093/bioinformatics/btq706

Shen W., Le S., Li Y., Hu F. 2016. SeqKit: A cross-platform and ultrafast toolkit for FASTA/Q file manipulation. PLOS ONE. 11: e0163962. 10.1371/journal.pone.0163962

Shimodaira H. 2002. An approximately unbiased test of phylogenetic tree selection. Systematic Biology. 51:492–508. 10.1080/10635150290069913

Simpson J.T., Wong K., Jackman S.D., Schein J.E., Jones S.J.M., Birol İ. 2009. ABySS: A parallel assembler for short read sequence data. Genome Research. 19:1117–1123. 10.1101/gr.089532.108

Slater G.S.C., Birney E. 2005. Automated generation of heuristics for biological sequence comparison. BMC Bioinformatics. 6:31. 10.1186/1471-2105-6-31

Smith A.B., Peterson K.J. 2002. Dating the time of origin of major clades: Molecular clocks and the fossil record. Annual Review of Earth and Planetary Sciences. 30:65–88. 10.1146/annurev.earth.30.091201.140057

Song F., Li H., Liu G.H., Wang W., James P., Colwell D.D., Tran A., Gong S., Cai W., Shao R. 2019. Mitochondrial genome fragmentation unites the parasitic lice of Eutherian mammals. Systematic Biology. 68:430–440. 10.1093/sysbio/syy062

Strimmer K., von Haeseler A. 1997. Likelihood-mapping: A simple method to visualize phylogenetic content of a sequence alignment. PNAS. 94:6815–6819. 10.1073/pnas.94.13.6815

Sweet A.D., Boyd B.M., Johnson K.P. 2016. Cophylogenetic patterns are uncorrelated between two lineages of parasites on the same hosts. Biological Journal of the Linnean Society. 118: 813–828. 10.1111/bij.12771

Sweet A.D., Johnson K.P. 2015. Patterns of diversification in small New World ground doves are consistent with major geologic events. The Auk. 132:300–312. 10.1642/AUK-14-193.1

Sweet A.D., Johnson K.P. 2018. The role of parasite dispersal in shaping a host–parasite system at multiple evolutionary scales. Molecular Ecology. 27:5104–5119. 10.1111/mec.14937

To T.-H., Jung M., Lycett S., Gascuel O. 2016. Fast dating using least-squares criteria and algorithms. Systematic Biology. 65:82–97. 10.1093/sysbio/syv068

de Vienne D.M., Refrégier G., López-Villavicencio M., Tellier A., Hood M.E., Giraud T. 2013. Coespeciation vs hos-shift speciation: methods for testing, evidence from natural associations and relation to coevolution. New Phytologist. 198:347–385. 10.1111/nph.12150

Villa S.M., Altuna J.C., Ruff J.S., Beach A.B., Mulvey L.I., Poole E.J., Campbell H.E., Johnson K.P., Shapiro M.D., Bush S.E., Clayton D.H. 2019. Rapid experimental evolution of reproductive isolation from a single natural population. PNAS. 116:13440–13445. 10.1073/pnas.1901247116

Vogwill T., Fenton A., Brockhurst M.A. 2008. The impact of parasite dispersal on antagonistic host--parasite coevolution. Journal of Evolutionary Biology. 21:1252–1258. 10.1111/j.1420-9101.2008.01574.x

Wang A.Y., Peng Y.Q., Harder L.D., Huang J.F., Yang D.R., Zhang D.Y., Liao W.J. 2019. The nature of interspecific interactions and co-diversification patterns, as illustrated by the fig microcosm. New Phytologist. 224:1304–1315. 10.1111/nph.16176

Weiblen G.D., Bush G.L. 2002. Speciation in fig pollinators and parasites. Molecular Ecology. 11:1573–1578. 10.1046/j.1365-294X.2002.01529.x

Wilson J.S., Forister M.L., Dyer L.A., O’Connor J.M., Burls K., Feldman C.R., Jaramillo M.A., Miller J.S., Rodríguez-Castañeda G., Tepe E.J., Whitfield J.B., Young B. 2012. Host conservatism, host shifts and diversification across three trophic levels in two Neotropical forests. Journal of Evolutionary Biology. 25:532–546. 10.1111/j.1420-9101.2011.02446.x

Yang Z. 2007. PAML 4: Phylogenetic analysis by maximum likelihood. Molecular Biology and Evolution. 24:1586–1591. 10.1093/molbev/msm088

Yang Z., Rannala B. 2006. Bayesian estimation of species divergence times under a molecular clock using multiple fossil calibrations with soft bounds. Molecular Biology and Evolution. 23:212–226. 10.1093/molbev/msj024

Zhang C., Rabiee M., Sayyari E., Mirarab S. 2018. ASTRAL-III: polynomial time species tree reconstruction from partially resolved gene trees. BMC Bioinformatics 2018 19:6. 19:15–30. 10.1186/s12859-018-2129-y

