## Supplementary Data for "Origin and diversification of a globally distributed group of parasitic feather lice"

### SUPPLEMENTARY TABLES

**Table S1 (Excel file).** Metadata for samples of dove body lice, landfowl lice, and outgroup taxa used for whole genome sequencing and subsequent assembly for phylogenetic analysis.

**Table S2.** Normalized Robinson-Foulds distances between pairs of phylogenetic trees estimated from alignments of nuclear genes from dove body lice and outgroup taxa.

| | **1** | **2** | **3** | **4** | **5** | **6** | **7** | **8** | **9** |
|---|---|---|---|---|---|---|---|---|---|
| **2** | 0.0787 | | | | | | | | |
| **3** | 0.0112 | 0.0899 | | | | | | | |
| **4** | 0.0337 | 0.0899 | 0.0449 | | | | | | |
| **5** | 0.0000 | 0.0787 | 0.0112 | 0.0337 | | | | | |
| **6** | 0.0337 | 0.0899 | 0.0449 | 0.0000 | 0.0337 | | | | |
| **7** | 0.0562 | 0.0337 | 0.0674 | 0.0674 | 0.0562 | 0.0674 | | | |
| **8** | 0.0112 | 0.0899 | 0.0000 | 0.0449 | 0.0112 | 0.0449 | 0.0674 | | |
| **9** | 0.0000 | 0.0787 | 0.0112 | 0.0337 | 0.0000 | 0.0337 | 0.0562 | 0.0112 | |
| **10** | 0.0337 | 0.0899 | 0.0449 | 0.0000 | 0.0337 | 0.0000 | 0.0674 | 0.0449 | 0.0337 |

1: Concatenated tree (max 60% missing data, all sites)
2: ASTRAL tree (max 90% missing data, 1+2 sites)
3: ASTRAL tree (max 90% missing data, all sites)
4: Concatenated tree (max 90% missing data, 1+2 sites)
5: Concatenated tree (max 90% missing data, all sites)
6: Concatenated tree (max 60% missing data, 1+2 sites)
7: ASTRAL tree (max 60% missing data, 1+2 sites)
8: ASTRAL tree (max 60% missing data, all sites)
9: Concatenated tree (no filtering, all sites)
10: Concatenated tree (no filtering, 1+2 sites)

**Table S3.** Results from Approximately Unbiased tests between nuclear phylogenies of dove body lice. All trees were compared to the most likely tree estimated from a concatenated alignment (60% max missing data). DeltaL indicates the difference in log-likelihood from the maximum.

| | LogL | DeltaL | p-AU |
|---|---|---|---|
| **1** | -61883684.9800 | 0.0001 | 0.4490 |
| **2** | -61894778.6000 | 11094.0000 | 0.000032 |
| **3** | -61892276.9100 | 8591.9000 | 1.55E-42 |
| **4** | -61884938.2400 | 1253.3000 | 0.0003 |
| **5** | -61883684.9800 | 0.0000 | 0.5150 |
| **6** | -61884938.2400 | 1253.3000 | 0.0004 |
| **7** | -61891621.5400 | 7936.6000 | 0.0003 |
| **8** | -61892276.9100 | 8591.9000 | 1.49E-42 |
| **9** | -61883684.9800 | 0.0003 | 0.4920 |
| **10** | -61884938.24 | 1253.3 | 0.0002 |

1: Concatenated tree (max 60% missing data, all sites)
2: ASTRAL tree (max 90% missing data, 1+2 sites)
3: ASTRAL tree (max 90% missing data, all sites)
4: Concatenated tree (max 90% missing data, 1+2 sites)
5: Concatenated tree (max 90% missing data, all sites)
6: Concatenated tree (max 60% missing data, 1+2 sites)
7: ASTRAL tree (max 60% missing data, 1+2 sites)
8: ASTRAL tree (max 60% missing data, all sites)
9: Concatenated tree (no filtering, all sites)
10: Concatenated tree (no filtering, 1+2 sites)

### SUPPLEMENTARY FIGURES

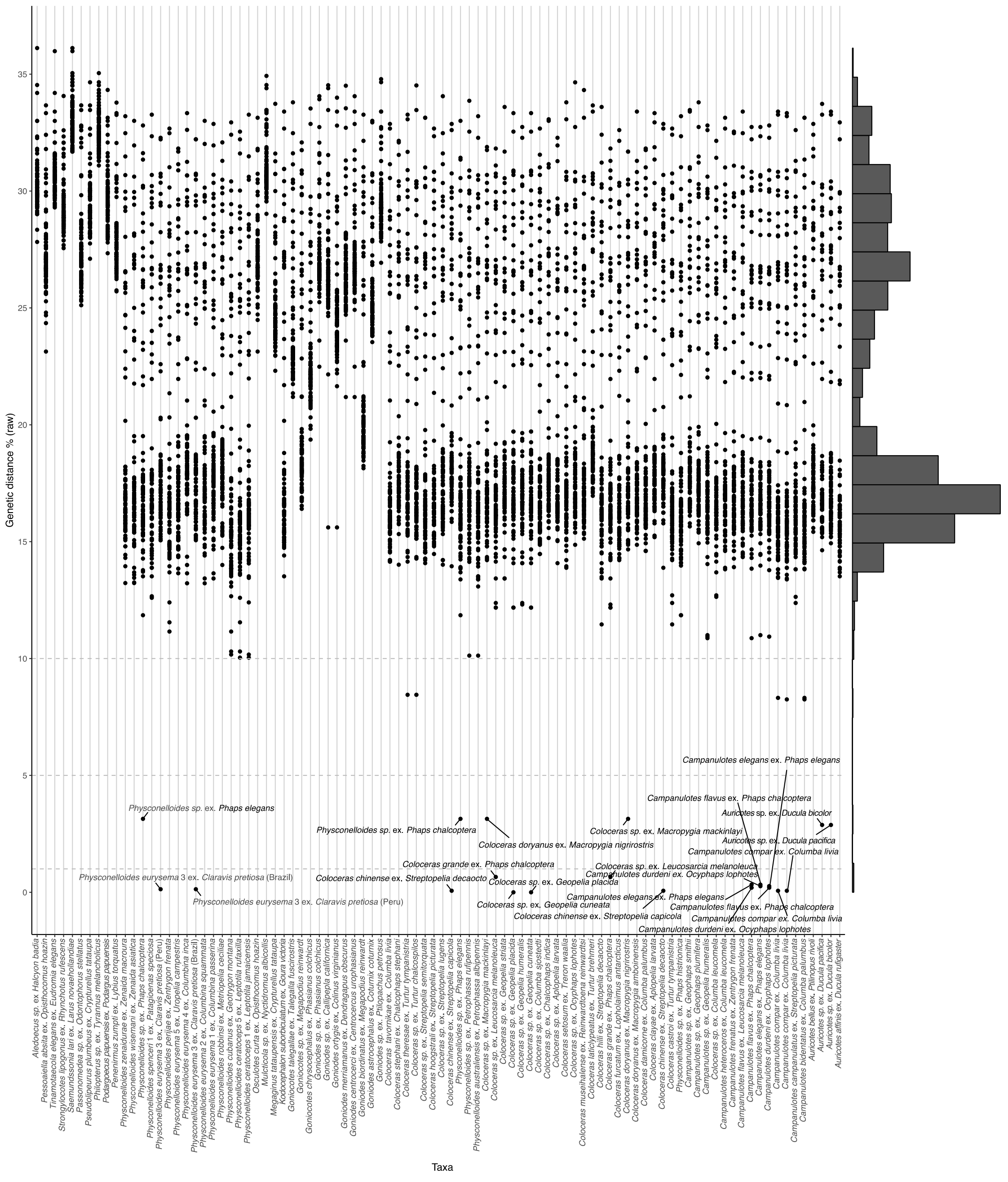


**Figure S1.** Uncorrected pairwise distances of mitochondrial *cox1* sequences from dove body lice. Taxa with distances below the 5% threshold for delimiting Operational Taxonomic Units (OTUs) are indicated with labels next to the respective points on the plot. These pairs of taxa were collapsed for subsequent analysis.

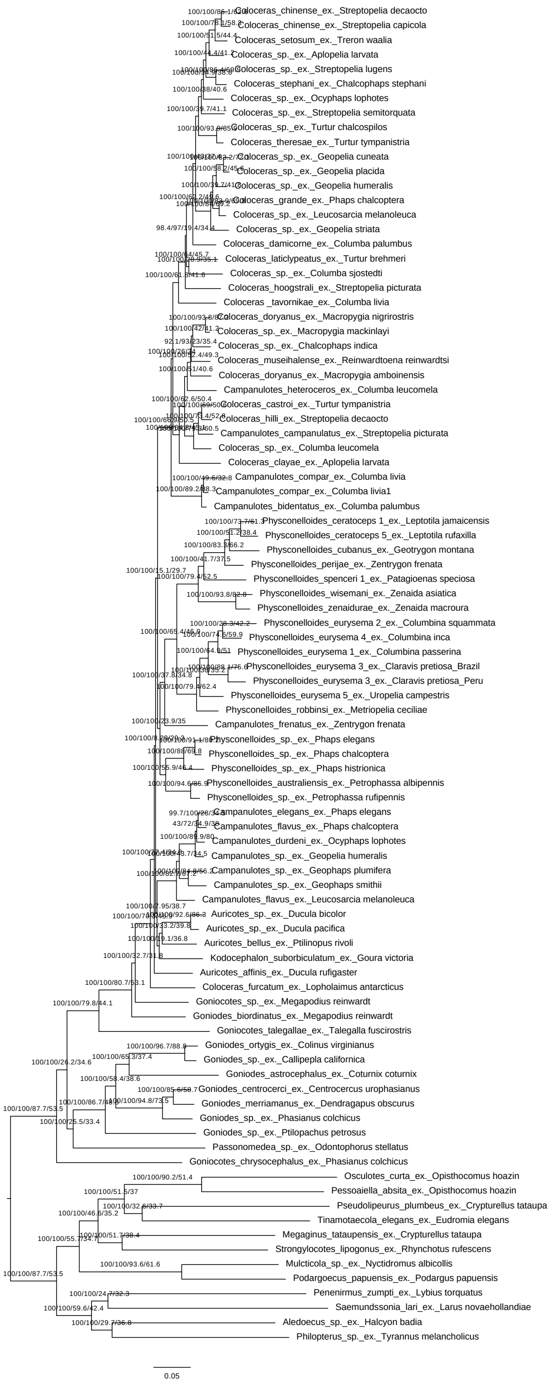


**Figure S1.** Maximum likelihood phylogeny of dove body lice, landfowl lice, and outgroup taxa based on a concatenated alignment of 2,367 orthologous genes with 60% maximum missing data (40% complete). Branch support values are shown above associated branches, and include SH-aLRT, bootstrap, gene concordance, and site concordance factor values, respectively. Branch lengths given in nucleotide substitutions per site.

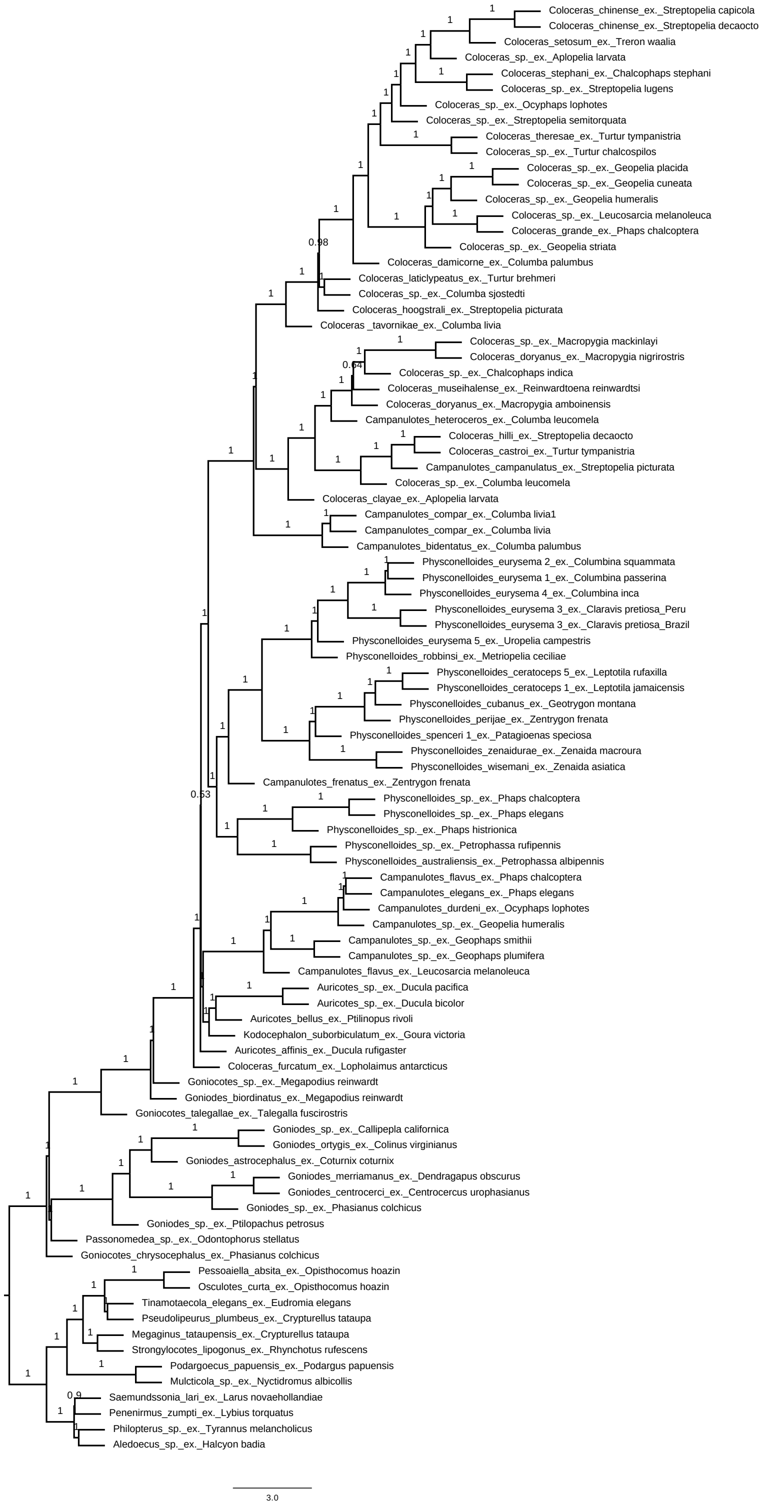


**Figure S2.** Species tree of dove body lice, landfowl lice, and outgroups taxa summarized from 2,367 genes trees estimated from alignments with 60% maximum missing data (40% complete). Local posterior probability support values are shown above associated branches. Internal branch lengths are in coalescent units. Branch lengths of terminal branches are not meaningful.

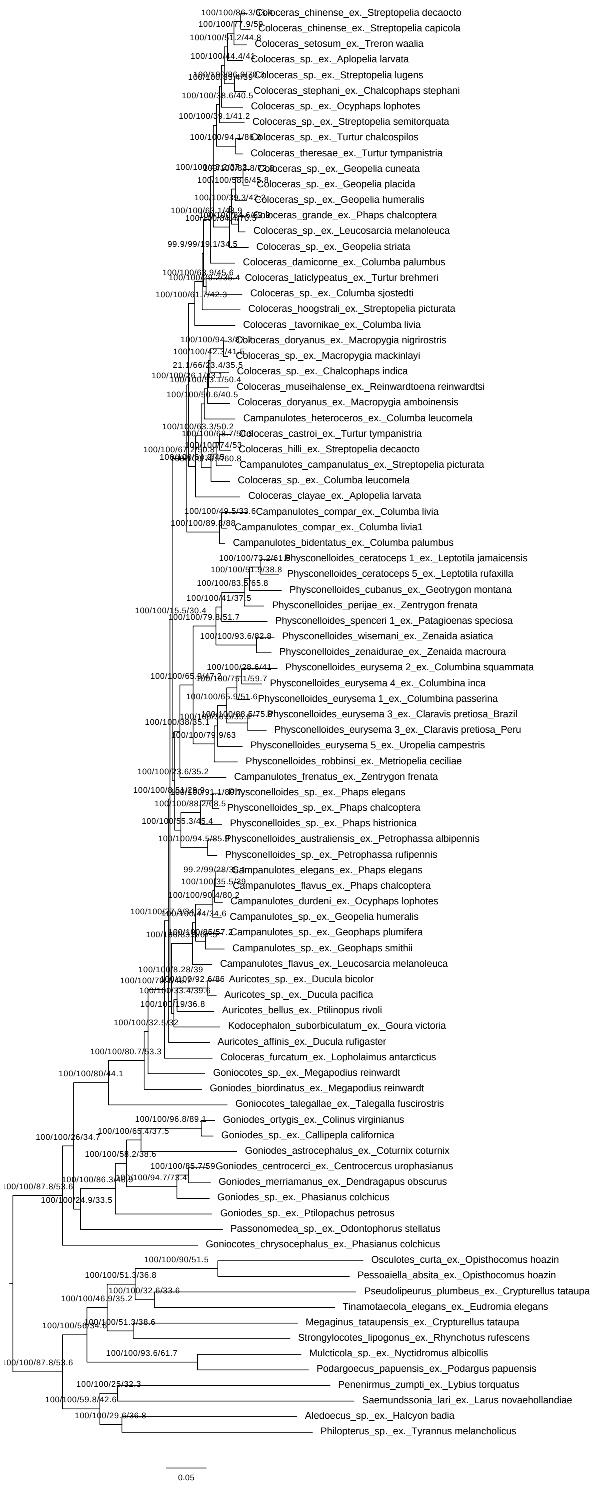

**Figure S3.** Maximum likelihood phylogeny of dove body lice, landfowl lice, and outgroups taxa based on a concatenated alignment of 2,367 orthologous genes with 10% maximum missing data (90% complete). Branch support values are shown above associated branches, and include SH-aLRT, bootstrap, gene concordance, and site concordance factor values, respectively.

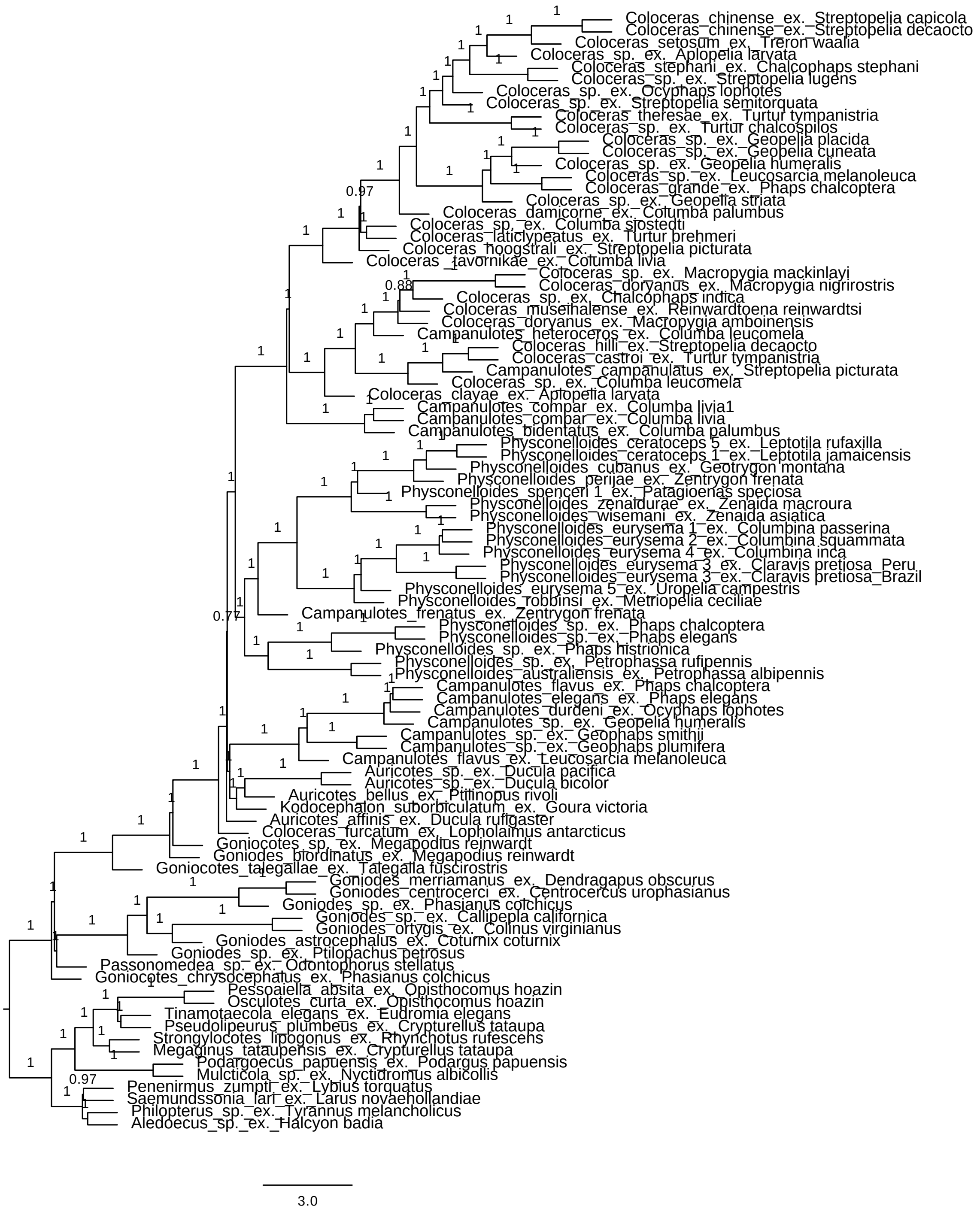


**Figure S4.** Species tree of dove body lice, landfowl lice, and outgroups taxa summarized from 2,367 genes trees estimated from alignments with 10% maximum missing data (90% complete). Local posterior probability support values are shown above associated branches. Internal branch lengths are in coalescent units. Branch lengths of terminal branches are not meaningful.

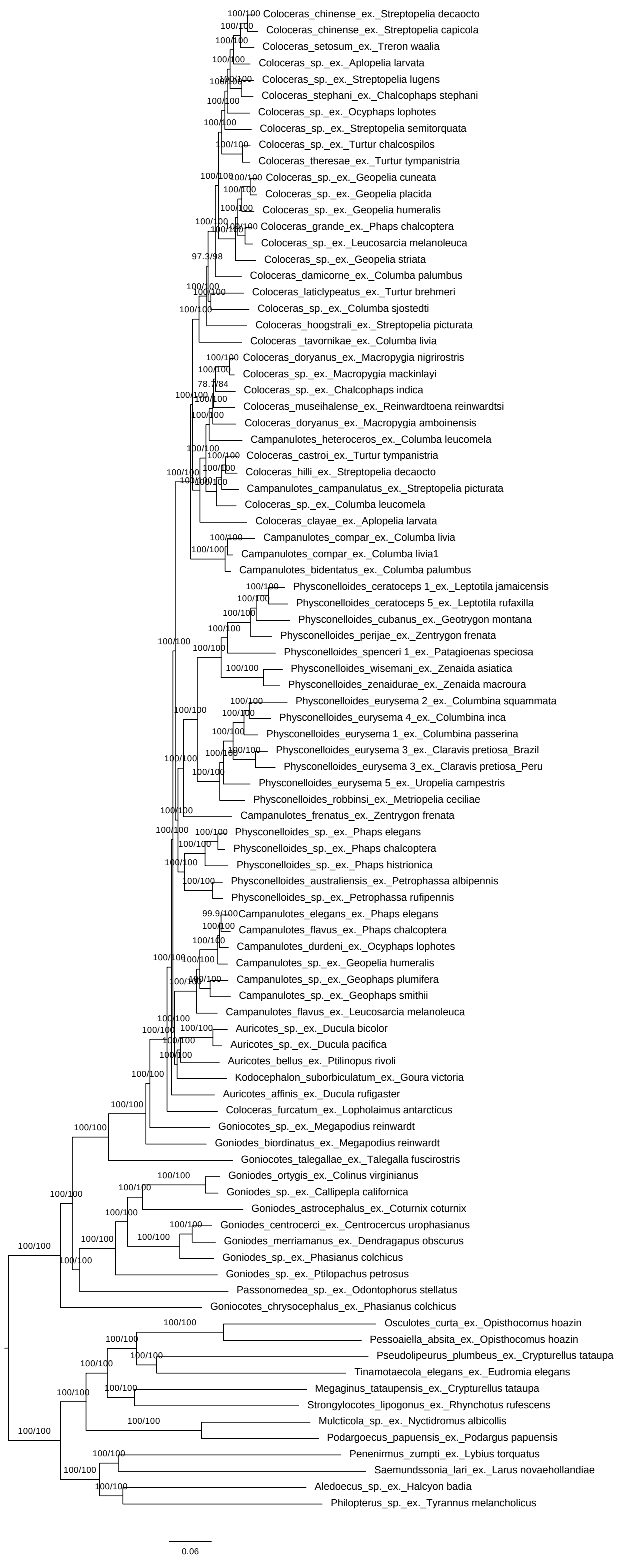


**Figure S5**. Maximum likelihood phylogeny of dove body lice, landfowl lice, and outgroups taxa based on a concatenated alignment of 2,367 orthologous genes with no filtering based on missing data (all sites). Branch support values are shown above associated branches, and include SH-aLRT and bootstrap values, respectively.

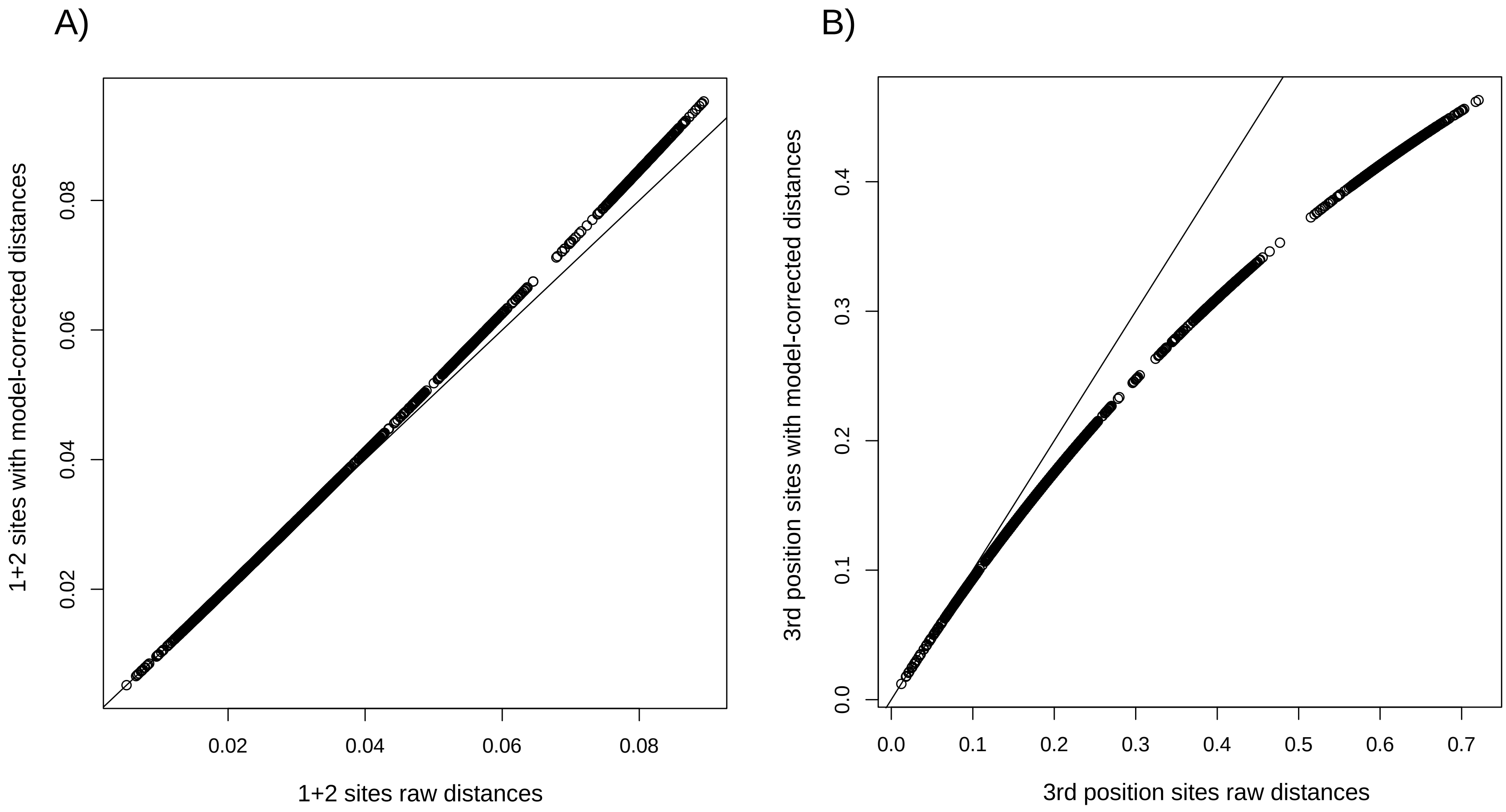


**Figure S6**. Saturation plots for a concatenated alignment of 2,367 nuclear genes from dove body lice (40% complete alignment).

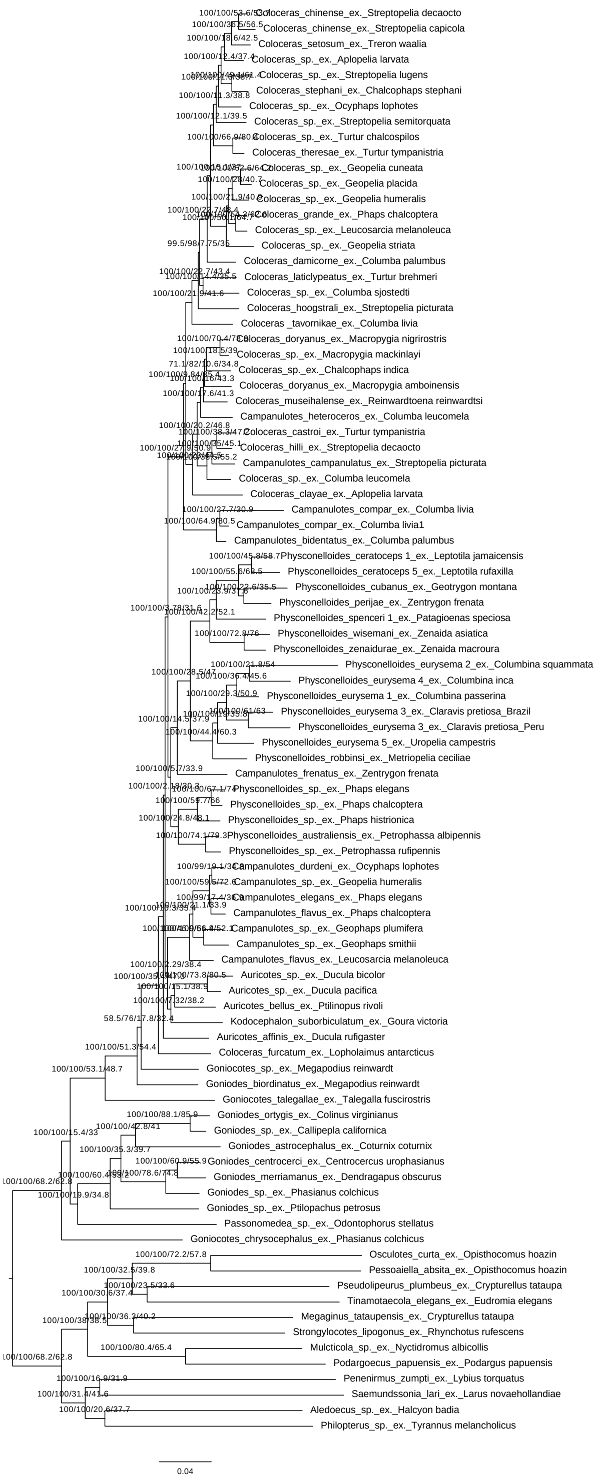

**Figure S7.** Maximum likelihood phylogeny of dove body lice, landfowl lice, and outgroups taxa based on a concatenated alignment of 2,367 orthologous genes with 60% maximum missing data (40% complete) and only first and second codon positions. Branch support values are shown above associated branches, and include SH-aLRT, bootstrap, gene concordance, and site concordance factor values, respectively.

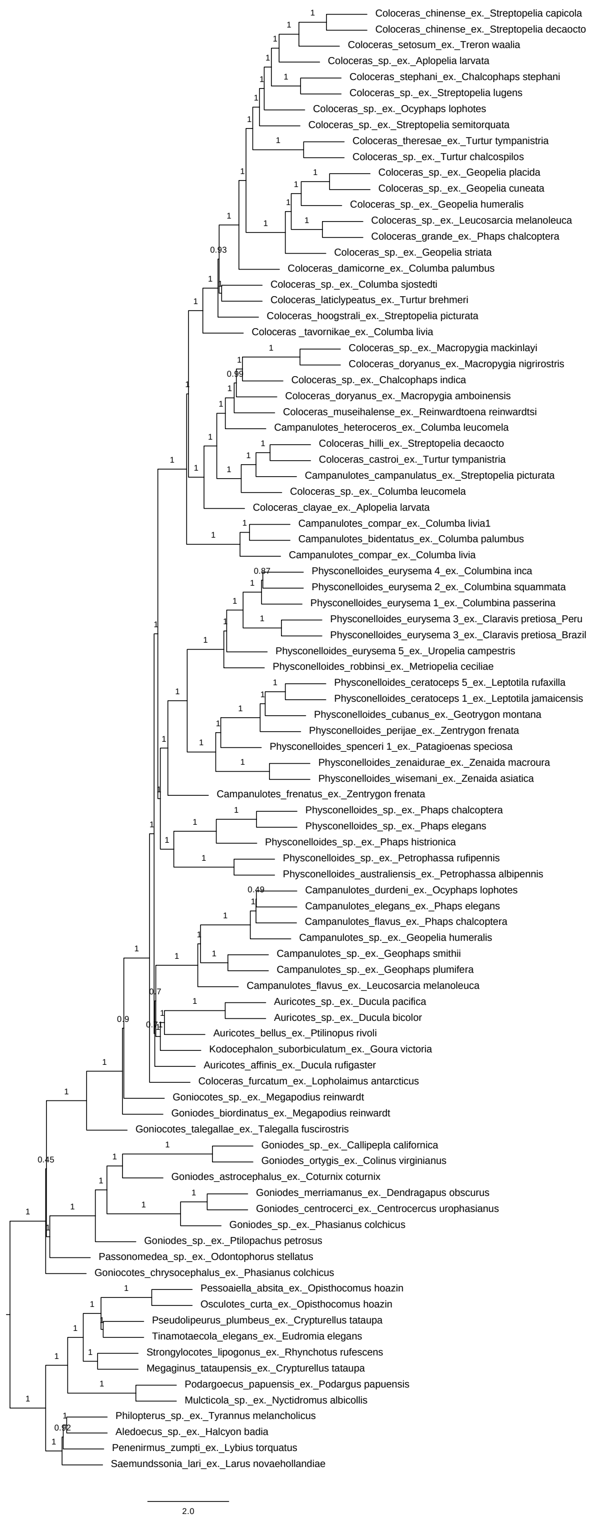


**Figure S8.** Species tree of dove body lice, landfowl lice, and outgroups taxa summarized from 2,367 genes trees estimated from alignments with 60% maximum missing data (40% complete) and only first and second codon positions. Local posterior probability support values are shown above associated branches. Internal branch lengths are in coalescent units. Branch lengths of terminal branches are not meaningful.

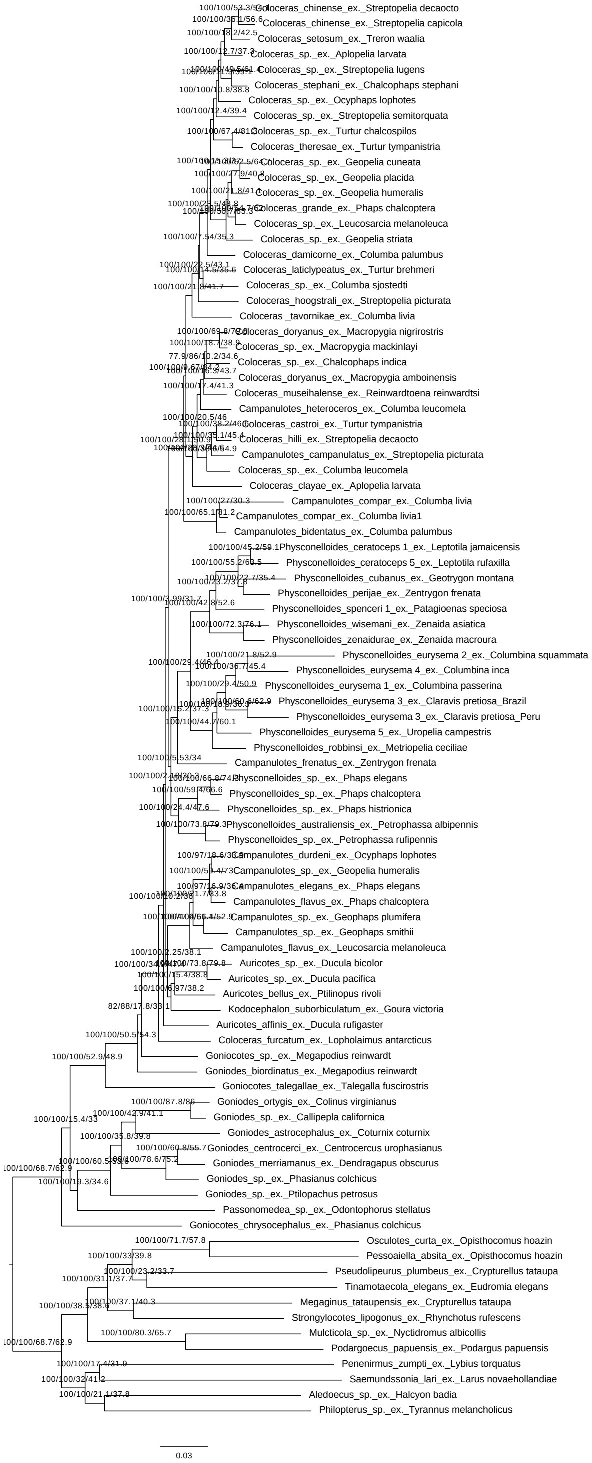


**Figure S9.** Maximum likelihood phylogeny of dove body lice, landfowl lice, and outgroups taxa based on a concatenated alignment of 2,367 orthologous genes with 10% maximum missing data (60% complete) and only first and second codon positions. Branch support values are shown above associated branches, and include SH-aLRT, bootstrap, gene concordance, and site concordance factor values, respectively.

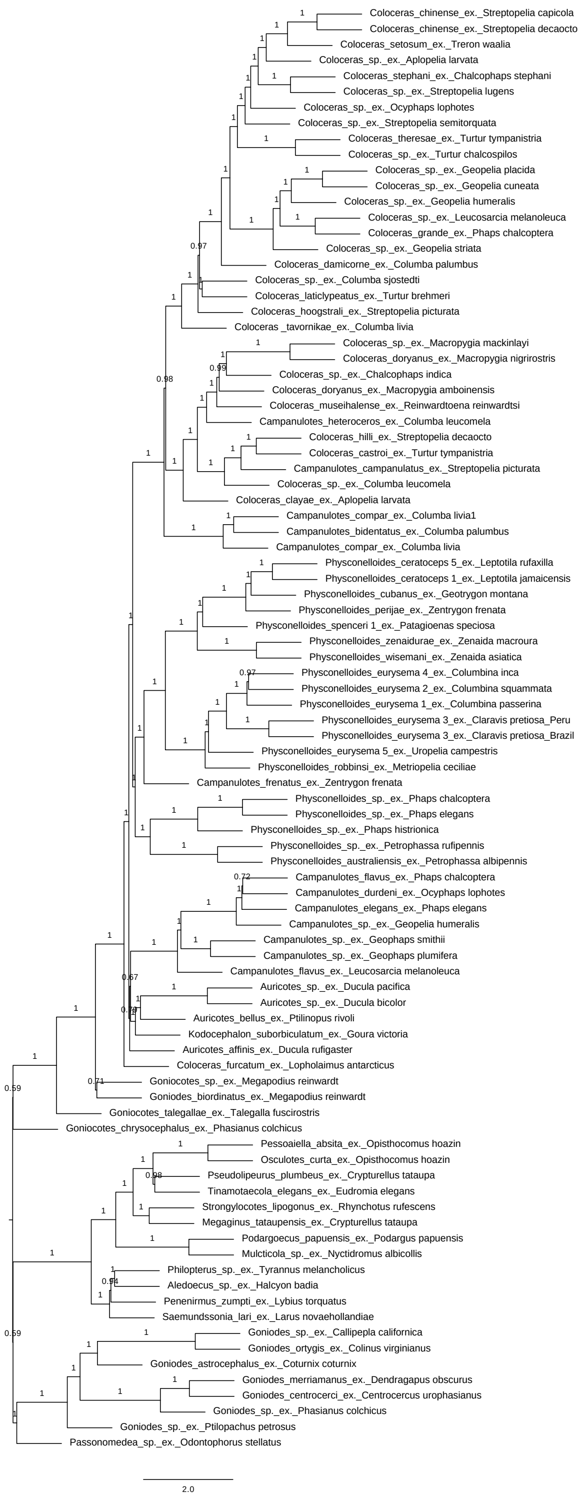


**Figure S10.** Species tree of dove body lice, landfowl lice, and outgroups taxa summarized from 2,367 genes trees estimated from alignments with 10% maximum missing data (90% complete) and only first and second codon positions. Local posterior probability support values are shown above associated branches. Internal branch lengths are in coalescent units. Branch lengths of terminal branches are not meaningful.

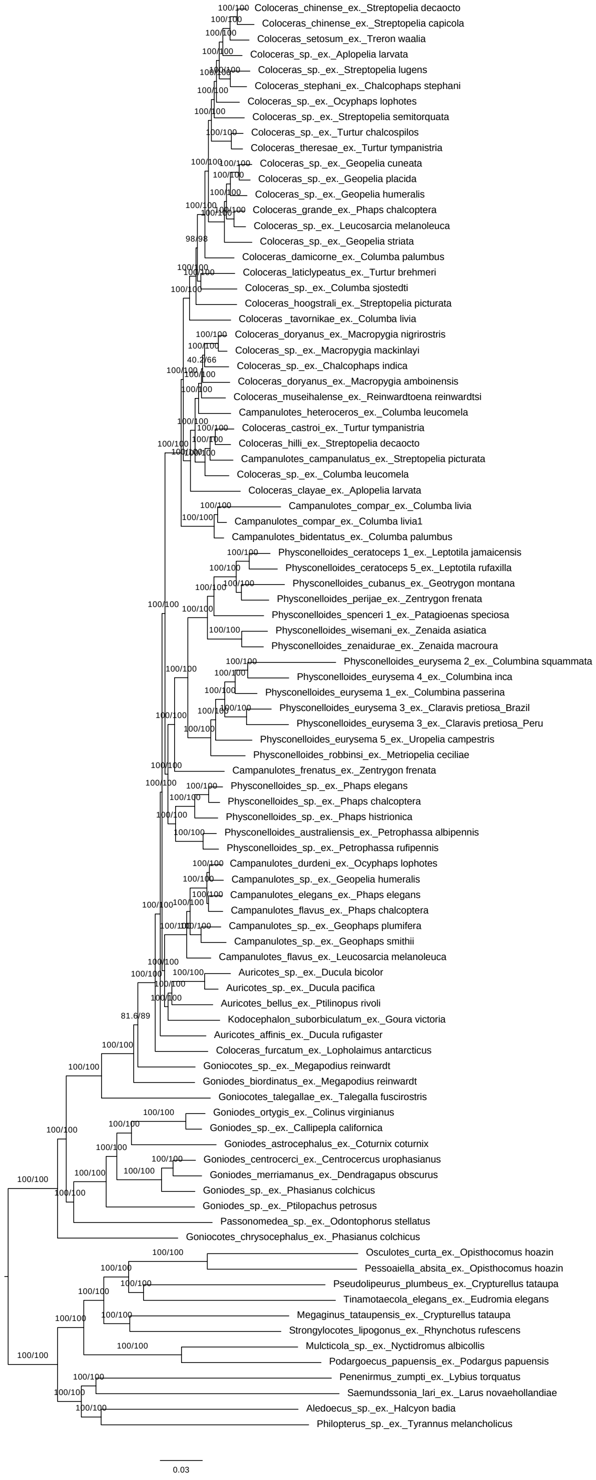


**Figure S11.** Maximum likelihood phylogeny of dove body lice, landfowl lice, and outgroups taxa based on first and second codon position sites in a concatenated alignment of 2,367 orthologous genes with no filtering based on missing data. Branch support values are shown above associated branches, and include SH-aLRT and bootstrap values, respectively.

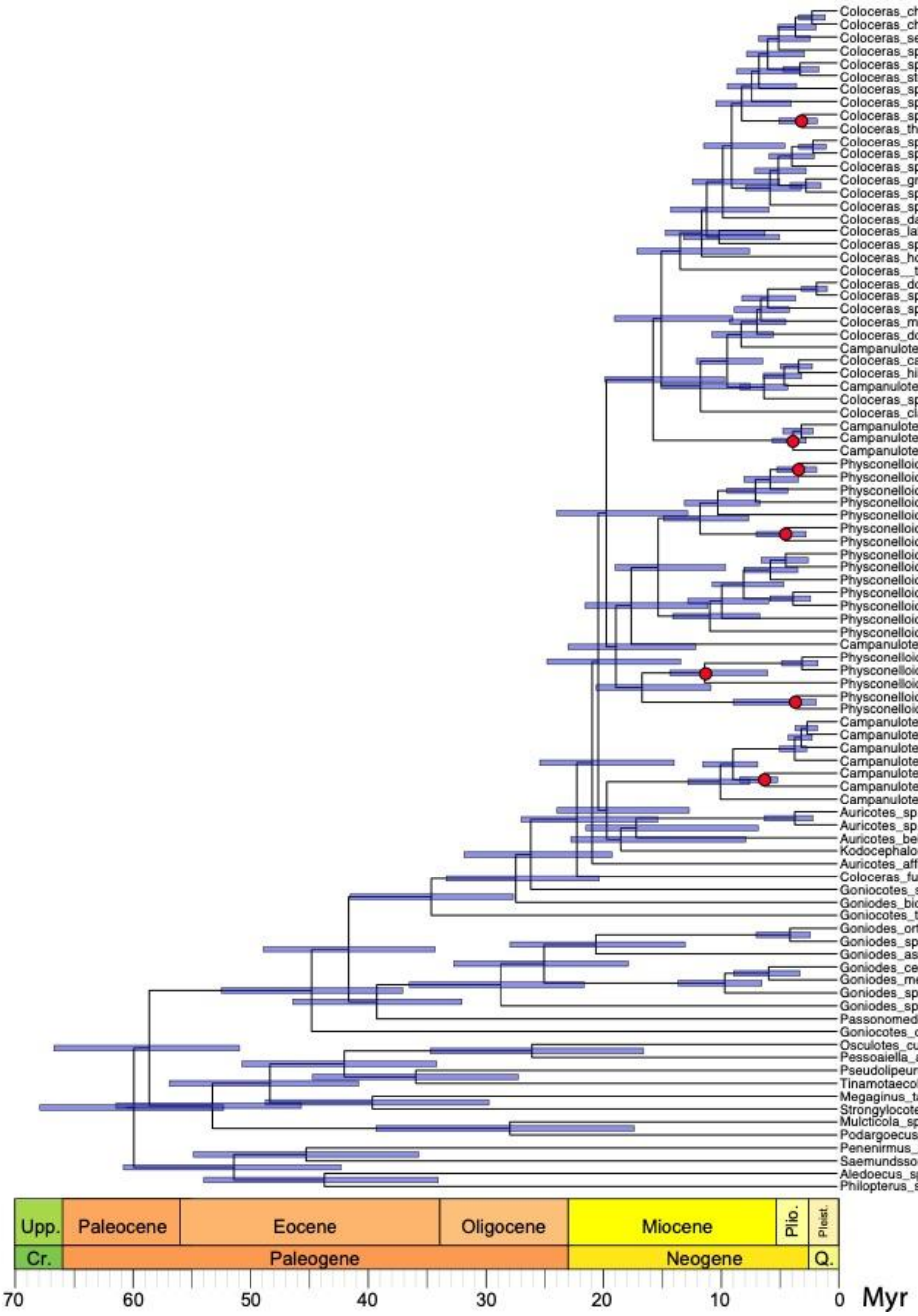


**Figure S12.** Dated phylogenetic tree of dove body lice, landfowl lice, and outgroups taxa from MCMCTree. 95% Highest Posterior Density (HPD) intervals are shown as blue bars at each node. Secondary calibration points inferred from host-parasite codivergence events are indicated with red circles.

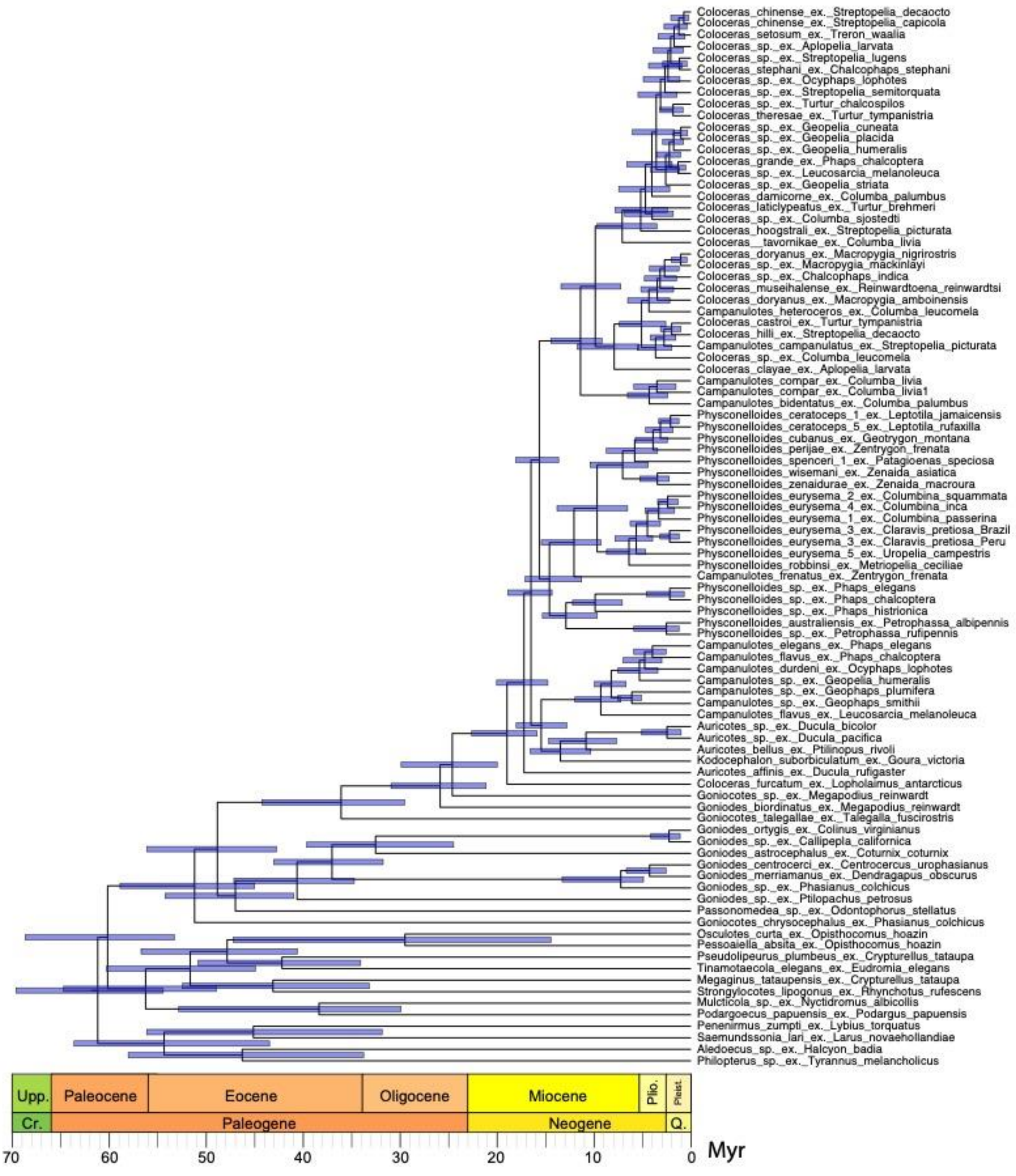


**Figure S13.** Dated phylogenetic tree of dove body lice, galliform lice, and outgroups taxa from a second run of MCMCTree. 95% Highest Posterior Density (HPD) intervals are shown as blue bars at each node.

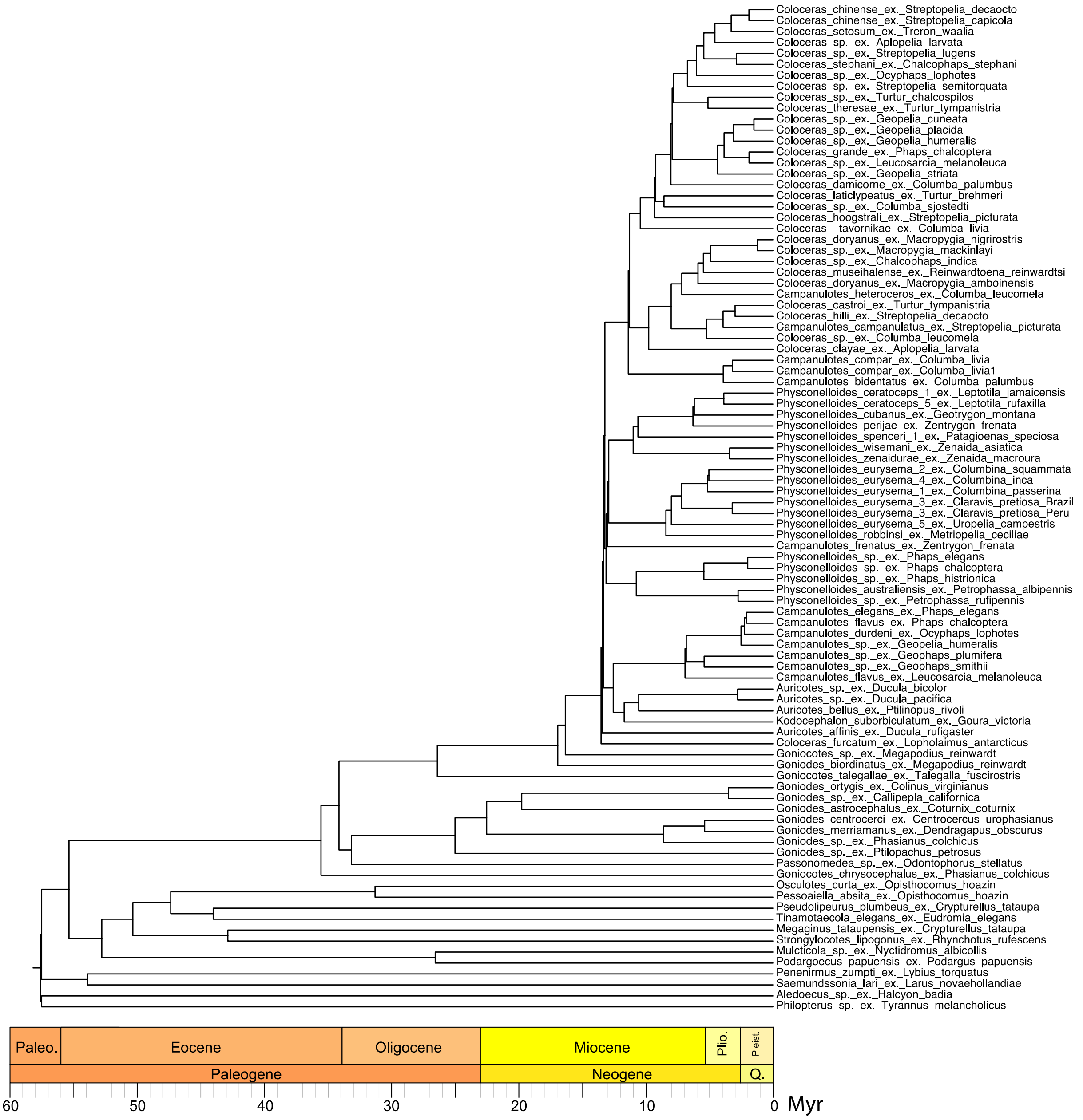


**Figure S14.** Dated phylogenetic tree of dove body lice, galliform lice, and outgroups taxa from the LSD2 method in IQTree2.

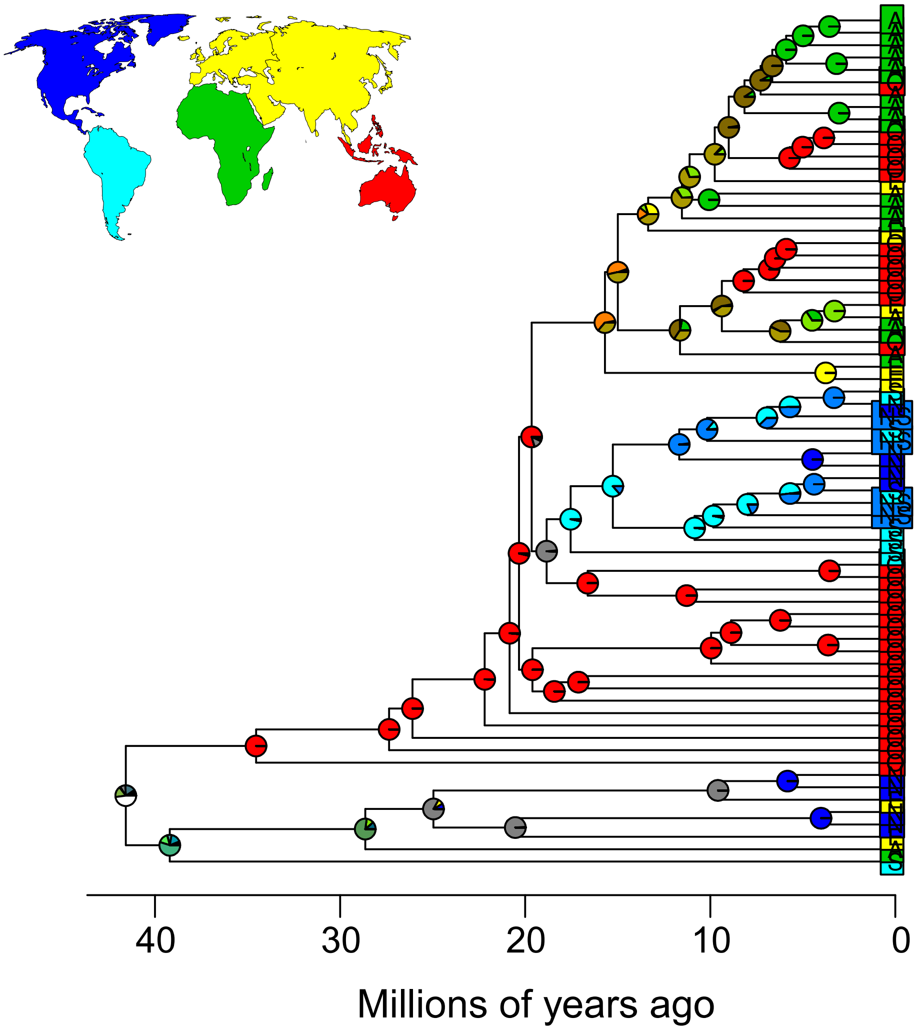


**Figure S15.** Dated phylogenetic tree of dove body lice and galliform lice under a DEC model in BioGeoBEARS. The DEC model was the best model with the jump (+J) parameters. Branch lengths are scaled to millions of years. Circles at the tips show the current known distributions of that taxon according to one of five biogeographic regions: Africa (green), Australasia (red), Eurasia (yellow), North America (dark blue), South America (light blue). Taxa found in both North and South America are shown with a medium-dark blue. Pie charts at each node indicate the likelihood the ancestor lived in a particular biogeographic region.

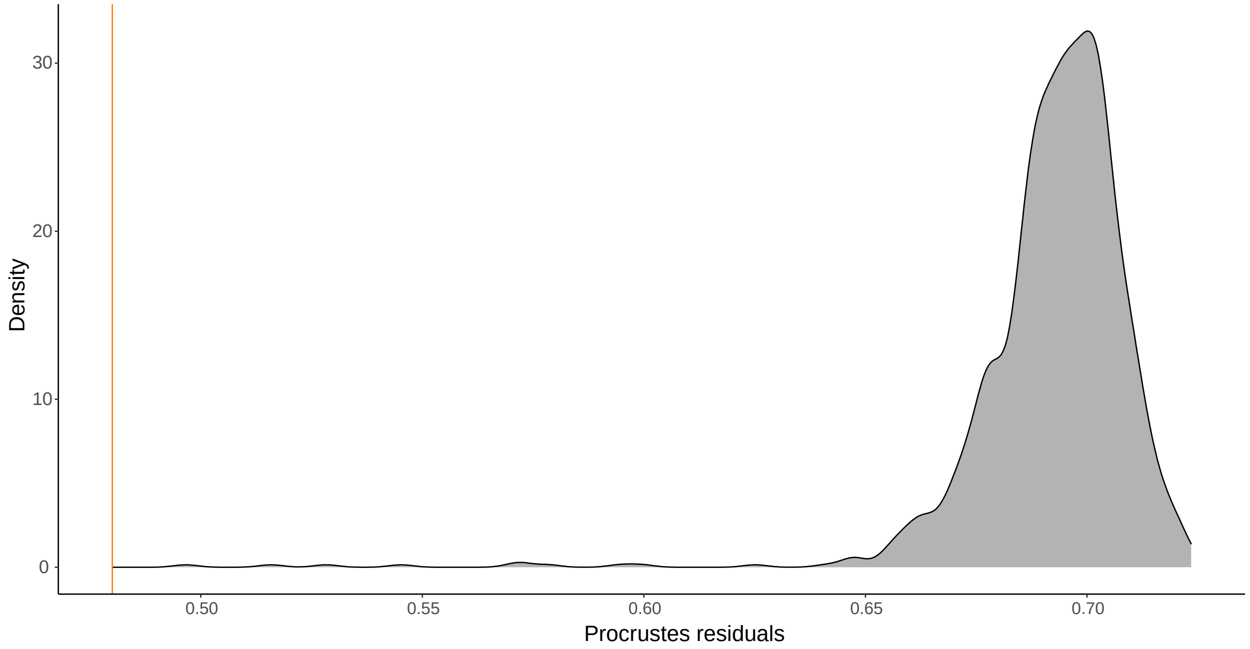


**Figure S16.** Distribution of Procrustes residuals from randomized associations in comparisons of dove wing louse and dove body louse phylogenies in PACo. The observed residual value is indicated with an orange vertical line.

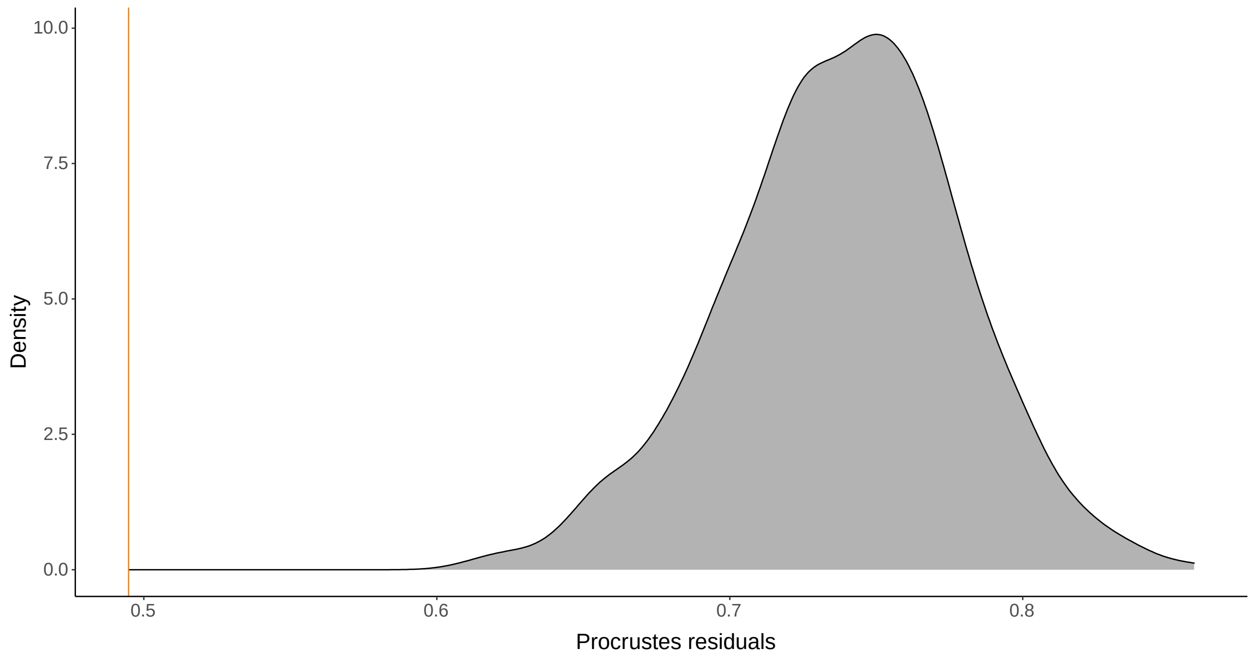


**Figure S17.** Distribution of Procrustes residuals from randomized phylogenies in comparisons of dove wing louse and dove body louse phylogenies in PACo. The observed residual value is indicated with an orange vertical line.

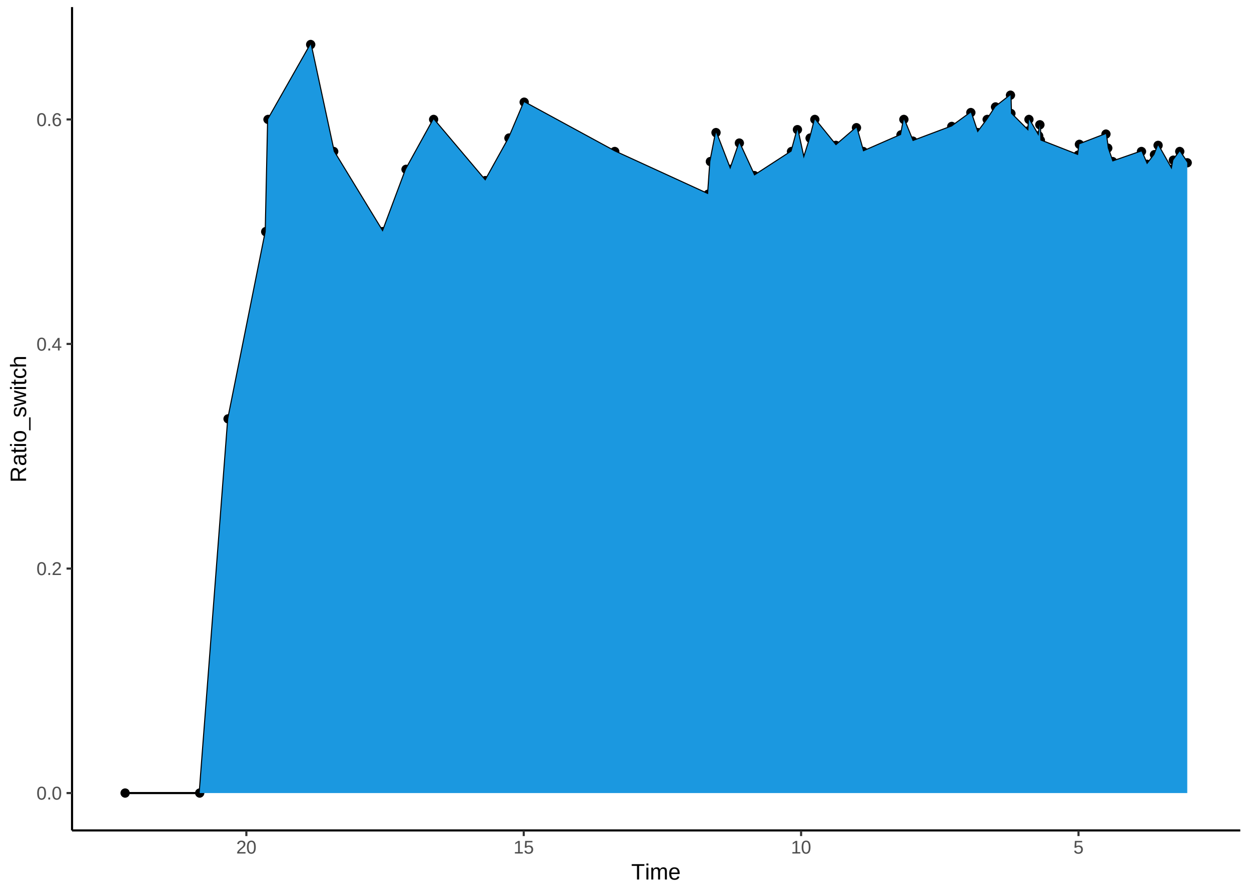


**Figure S18.** Proportion of cumulative speciation events in dove body lice that are attributed to host switches versus cospeciation events over time. X-axis indicates binds of time in millions of years before present.

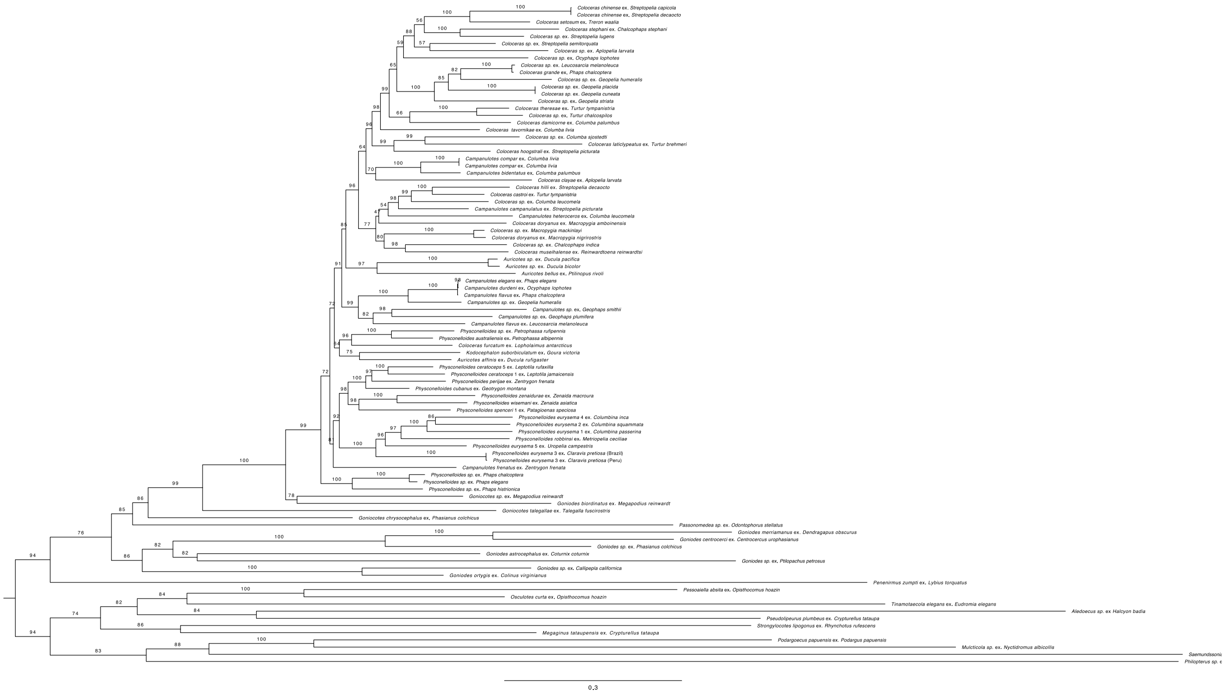

**Figure S19.** Maximum likelihood phylogeny of dove body lice, landfowl lice, and outgroup taxa based on the mitochondrial *cox1* gene. Bootstrap support values are shown above each branch. Branch lengths given in nucleotide substitutions per site.
